# L-type Ca^2+^ channels mediate regulation of glutamate release by subthreshold potential changes

**DOI:** 10.1101/2023.01.18.524500

**Authors:** Byoung Ju Lee, Unghwi Lee, Seung Hyun Ryu, Sukmin Han, Seung Yeon Lee, Jae Sung Lee, Anes Ju, Sunghoe Chang, Suk-Ho Lee, Sung Hyun Kim, Won-Kyung Ho

## Abstract

Subthreshold depolarization enhances neurotransmitter release evoked by action potentials and plays a key role in modulating synaptic transmission by combining analog and digital signals. This process is known to be Ca^2+^-dependent. However, the underlying mechanism of how small changes in basal Ca^2+^ caused by subthreshold depolarization can regulate transmitter release triggered by a large increase in local Ca^2+^ is not well understood. This study aimed to investigate the source and signaling mechanisms of Ca^2+^ that couple subthreshold depolarization with the enhancement of glutamate release in hippocampal cultures and CA3 pyramidal neurons. Subthreshold depolarization increased presynaptic Ca^2+^ levels, the frequency of spontaneous release, and the amplitude of evoked release, all of which were abolished by blocking L-type Ca^2+^ channels. A high concentration of intracellular Ca^2+^ buffer or blockade of calmodulin and phospholipase C abolished depolarization induced increases in transmitter release. Estimation of the readily releasable pool size using hypertonic sucrose showed depolarization induced increases in readily releasable pool size, and this increase was abolished by blockade of calmodulin or phospholipase C. Our results provide mechanistic insights into the modulation of transmitter release by subthreshold potential change and highlight the role of L-type Ca^2+^ channels in coupling subthreshold depolarization to the activation of Ca^2+^-dependent signaling molecules that regulate transmitter release.

**SIGNIFICANCE:** Neuronal activities are encoded by action potentials, but subthreshold changes in resting membrane potentials also play important roles in regulating neuronal functions including synaptic transmission. It is, however, poorly understood how small changes in basal Ca^2+^ induced by subthreshold depolarization regulate transmitter release triggered by a large increase in local Ca^2+^ in presynaptic terminals. We demonstrate that L-type Ca^2+^ channels are the major source of presynaptic Ca^2+^ influx at basal state and during subthreshold depolarization, resulting in the activation of signaling molecules such as calmodulin and phospholipase C, which facilitate transmitter release by increasing both release probability and the readily releasable pool size. Our results provide mechanistic insight into how subthreshold potential changes contribute to regulating transmitter release.

## INTRODUCTION

Synaptic transmission, a core process of information flow mediated by neurotransmitter release from presynaptic terminals, is triggered by action potentials (APs). Although APs are generally considered all-or-none signals, they are modulated by subthreshold potential changes (1), which in turn affect synaptic strength (2, 3). In addition, subthreshold changes in somatic potentials can electrotonically spread to axon terminals and modulate spontaneous or asynchronous release (3). The involvement of subthreshold depolarization in the emergence of place field spiking (4) and the propensity to have place fields (5) has also been highlighted in recent studies. Considering that the resting membrane potential (RMP) is not fixed but fluctuates in the subthreshold range by the alteration of extracellular K^+^ concentrations, ion channel activities, and synaptic activities, the RMP is a key player in the analog-digital modulation of synaptic transmission and neural activities (6). However, the mechanisms underlying the modulation of synaptic transmission and neural activity by RMP changes are not well understood.

Neurotransmitter release at synapse is orchestrated by numerous molecular mechanisms that govern multiple steps of synaptic vesicle dynamics, including vesicle priming, fusion, and recycling (7). It is well known that the low-affinity Ca^2+^ sensor synaptotagmins transduce a large increase in presynaptic Ca^2+^ into the fusion-pore opening of synaptic vesicles for transmitter release (8). Therefore, an AP-induced large increase in Ca^2+^ that persists for a short period is the initial step that triggers transmitter release. Interestingly, the slow dynamics of Ca^2+^ changes in a lower concentration range can regulate Ca^2+^-triggered transmitter release by activating numerous Ca^2+^-dependent signaling molecules. Calmodulin (CaM) plays a key role in vesicle priming and replenishment of the vesicle pool, thereby regulating short-term plasticity (9, 10). Diacylglycerol (DAG), a product of phospholipase C (PLC), increases transmitter release possibly by increasing fusion willingness (11, 12). Munc13 proteins (mammalian homologs of *Caenorhabditis elegans* UNC13), which are essential regulators of synaptic vesicle priming (13, 14), have binding sites for both CaM and DAG (9, 15, 16), suggesting a possibility that their function is regulated by these signaling molecules. The functional importance of Ca^2+^-dependent signals in synaptic transmission has been mostly investigated to understand the short-term plasticity mechanism induced by high-frequency activity (17). It is of interest to determine whether these signaling cascades also contribute to basal synaptic transmission regulated by subthreshold potential changes.

The role of presynaptic Ca^2+^ currents, including P/Q-, N-, and R-type Ca^2+^ currents mediated by Ca_V_2.1, Ca_V_2.2, and Ca_V_2.3 channels, respectively, in Ca^2+^-triggered transmitter release, is well established (18–20). However, it has been reported that L-type Ca^2+^ currents (LTCCs) do not participate in neurotransmitter release in most neurons (21, 22), except inner hair cells (23, 24) and bipolar cells in retina (25, 26). However, increased GABA release by increasing L-type Ca^2+^ currents using Bay K 8644 was recently reported in cerebellar molecular layer interneurons (27), suggesting the contribution of LTCCs to synaptic transmission. Considering that substitution of the synaptic protein interacting (synprint) site from Ca_V_2.1 channels in Ca_V_1.2 was sufficient to establish synaptic transmission initiated by LTCCs (28), the localization of Ca^2+^ channels at synaptic sites is important for their role in synaptic transmission. A recent study showed that chronic treatment with lipopolysaccharide increased Ca_V_1.2 channels at excitatory presynaptic terminals and their contribution to the increase of glutamate release (29), suggesting the possibility that Ca_V_1.2 localization and its contribution to synaptic transmission can be regulated. It would be intriguing to investigate whether LTCC-mediated Ca^2+^ influx plays a role in regulating neurotransmitter release under physiological conditions, such as subthreshold depolarization.

In the present study, we found that LTCCs played a key role in the enhancement of transmitter release by subthreshold depolarization. Furthermore, we demonstrated that the depolarization induced increase in transmitter release is mediated by increased readily releasable pool (RRP) size, which is attributable to CaM and PLC activation by LTCC-dependent elevation of presynaptic Ca^2+^ levels. Our results provide mechanistic insight into how RMP changes in the subthreshold range contribute to regulating transmitter release.

## RESULTS

### LTCCs regulate both spontaneous and evoked glutamate release

To test the contribution of L-type calcium channels (LTCCs) at presynaptic terminals to neurotransmitter release, we used autaptic cultured hippocampal neurons that enables manipulation of the presynaptic compartment environment. In the control condition, whole-cell voltage-clamp recordings were performed in aCSF containing 2 mM Ca^2+^ and 2.5 mM K^+^ with a pipette solution containing 0.1 mM ethylene glycol-bis (β-aminoethyl ether)-N,N,N’,N’-tetraacetic acid (EGTA). Excitatory neurons were distinguished from inhibitory neurons by fast decay kinetics of synaptic currents (ranged from 3.4 to 13.8 ms with mean value of 6.9 ± 0.2 ms, supplementary Fig. 1, *N* = 94) as previously described (30). Miniature excitatory postsynaptic currents (mEPSCs) were recorded at the holding potential (HP) of −70 mV, while evoked excitatory postsynaptic currents (eEPSCs) were recorded by applying 2 ms depolarizing step pulses to 0 mV from the HP every 20 s. The mEPSC frequency and eEPSC amplitude measured under the experimental conditions were normalized to the control level. To assess the role of LTCCs in glutamate release, we examined the effects of drugs that inhibit or activate LTCCs on the mEPSC frequency or eEPSC amplitude. Nimodipine (Nimo, 10 μM), a blocker of LTCCs, significantly decreased mEPSC frequency (Fig. 1A1 and B, blue, 0.81 ± 0.01, *N* = 17, normalized to the control value) without affecting mEPSC amplitudes (Fig. 1C), whereas LTCC activator (10 μM Bay K 8644, Bay K) induced a significant increase in mEPSC frequency (Fig. 1A2 and B, green, 1.79 ± 0.11, *N* = 7, normalized to control). Nimo and Bay K also altered the eEPSC amplitude in the same direction as the mEPSC frequency (Figs. 1D and E). Nimo and Bay K decreased and increased the eEPSC amplitude, respectively. In addition, the paired-pulse ratio (PPR) was increased by Nimo and decreased by Bay K application (Fig. 1F), suggesting that presynaptic mechanisms are involved in the LTCCs’ effects on eEPSCs. To further confirm that the effects of Nimo and Bay K were specifically mediated by LTCCs, we prepared Ca_V_1.2 and Ca_V_1.3 double knockdown autapses (See Supplementary Materials and Methods), since LTCCs are mainly comprised of Ca_V_1.2 and Ca_V_1.3 subunits in hippocampus (31), and tested the effects of Nimo and Bay K. Neither the eEPSC amplitude nor the mEPSC frequency was affected by Nimo or Bay K (Fig. 1G-I and supplementary Fig. 2). These results showed that LTCCs contribute to evoked as well as spontaneous glutamate release at the physiological RMP.

**Figure 1.**
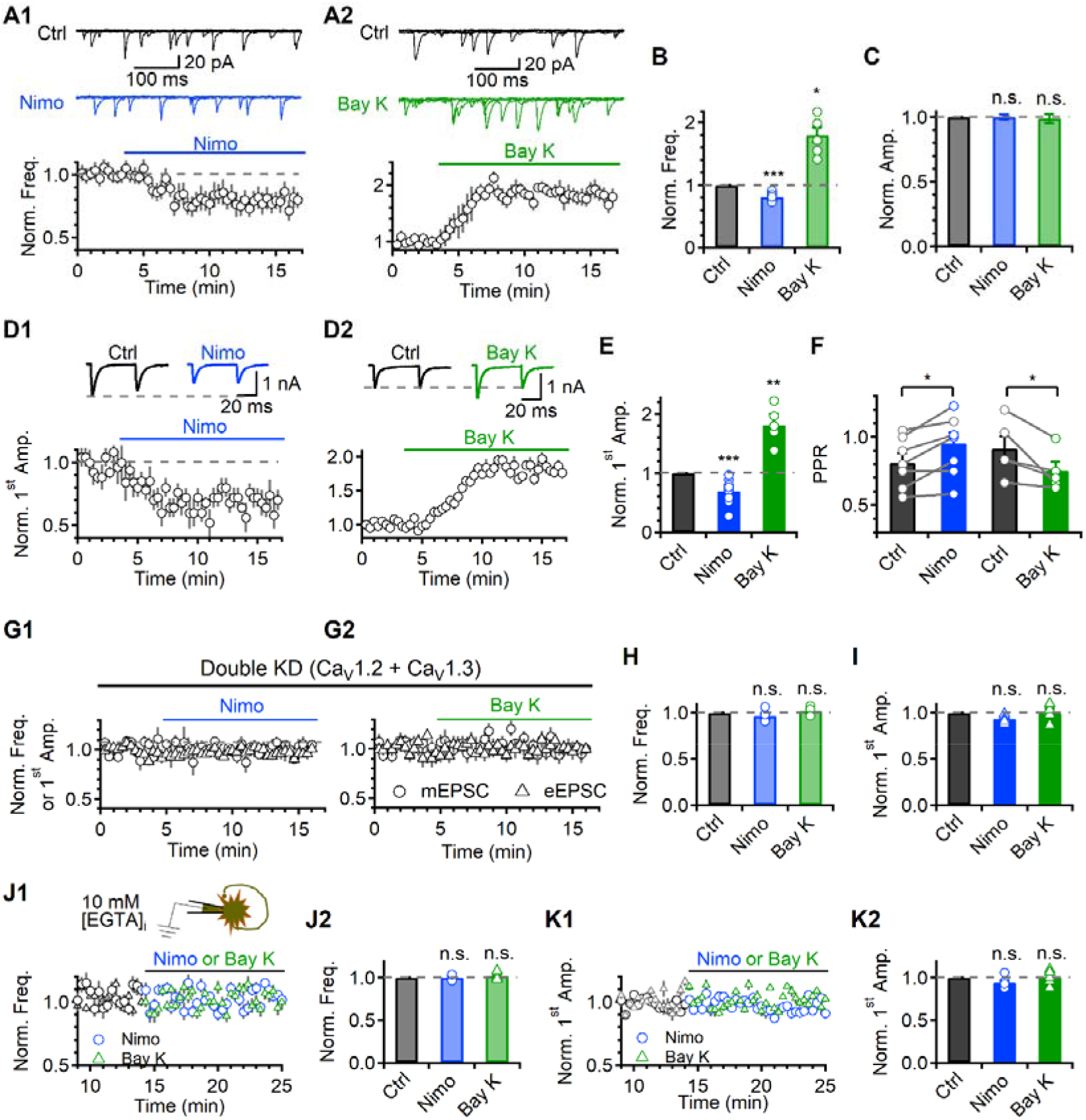
LTCCs regulate both spontaneous and evoked glutamate release. (A) Top. There are representative traces of mEPSCs in control, Nimo (A1, blue), and Bay K (A2, green). Five 500 ms long mEPSC traces were overlaid. Bottom. Average time courses of the normalized mEPSC frequency. The data were normalized by the mean mEPSC frequency of control. A dashed grey line indicates the control level. (B) A bar graph of average values of the normalized mEPSC frequency in different conditions. (C) A bar graph of the average values of the normalized mEPSC amplitude. (D) Top. There are representative traces of eEPSCs in the control, Nimo (D1), and Bay K (D2) conditions. The grey dashed line indicates the control first eEPSC peak amplitude. Bottom. Average time courses of the normalized first eEPSC amplitude. (E) A bar graph of the average values of the normalized first eEPSC amplitude in different conditions, compared to control. (F) A bar graph of the average values of the PPR in different conditions. (G) Average time courses of the normalized mEPSC frequency (circle) and first eEPSC amplitude (triangle) in Ca_V_1.2 and Ca_V_1.3 double knockdown autapses in presence of Nimo (G1) and Bay K (G2). (H) A bar graph of the average values of the normalized mEPSC frequency in different conditions, compared to control. (I) A bar graph of the average values of the normalized first eEPSC amplitude in different conditions, compared to control. (J1) Top. A schematic image of autaptic pyramidal neuron containing 10 mM EGTA internal patch pipette solution. Bottom. Average time courses of the normalized mEPSC frequency (Nimo, circle; Bay K, triangle). (J2) A bar graph of the average values of the normalized mEPSC frequency in different conditions, compared to control. (K1) Average time courses of the normalized first eEPSC amplitude. (K2) A bar graph of the average values of the normalized first eEPSC amplitude in different conditions, compared to control. The individual raw values are described in table S1.

In addition, we interestingly found that the effects of Nimo and Bay K on mEPSC frequency (Fig. 1J and supplementary Fig. 3B) and eEPSC amplitude (Fig. 1K and supplementary Fig. 3C) were completely abolished by 10 mM EGTA in the pipette solution, which inhibited global Ca^2+^ changes without affecting local Ca^2+^ increases near Ca^2+^ channels upon Ca^2+^ channel opening (32). The effect of 10 mM EGTA on the LTCC contribution differed from its effect on P/Q-, N-, and R-type contributions. Intracellular 10 mM EGTA did not affect the effects of P/Q-, N-, or R-type blockers (0.1 μM ω-agatoxin-IVA, 0.1 μM ω-conotoxin GVIA, and 100 μM NiCl_2_, respectively) on mEPSCs (supplementary Fig. 4), which is consistent with the notion that the contribution of P/Q-, N-, and R-type Ca^2+^ channels is mediated by local Ca^2+^ increases (30). There was no additive effect of Bay K on mEPSC frequency and eEPSC amplitude when all P/Q-, N-, and R-type VGCCs were blocked by the mixture of 0.1 μM Aga, 0.1 μM Cono, and 100 μM NiCl_2_ (3-mix, supplementary Fig. 5), implying that Ca^2+^ influx via LTCCs does not directly trigger exocytosis by increasing local Ca^2+^ near primed vesicles with nanodomain or microdomain coupling but indirectly augments vesicle release triggered by P/Q-, N-, and R-type VGCCs.

### V_m_-dependent regulation of spontaneous release is mediated by LTCCs-dependent changes in basal Ca^2+^

Several studies have reported the enhancement of neurotransmitter release by subthreshold depolarization (2, 3, 33), but the underlying mechanism remains unclear. We investigated whether membrane potential (V_m_) changes in the subthreshold range affect the glutamate release and, if so, whether LTCCs are involved in this mechanism. Autaptic cultured neurons allowed us to assess the relationship between V_m_ and spontaneous release by manipulating V_m_ in two ways: shifting the HP of patched neurons or changing the external K^+^ concentration ([K^+^]_e_). Lowering HP from −70 mV to −80 mV reduced mEPSC frequency (Fig. 2A1 and B, 0.82 ± 0.02, *N* = 12), while elevating HP from −70 mV to −60 mV increased mEPSC frequency (Fig. 2A1 and B, 1.49 ± 0.06, *N* = 11), indicating that V_m_ around the RMP dynamically impacts spontaneous glutamate release. Reduction of mEPSC amplitude by depolarization was detected (Supplementary Fig. 6, −80 mV, 17.66 ± 0.68; −70 mV, 15.98 ± 0.54; −60 mV, 14.2 ± 0.56 pA), which possibly reflect decreased driving force for non-selective cation currents. V_m_-dependent changes in mEPSC frequency were similarly observed when V_m_ was altered by changing the RMP with different [K^+^]_e_ (Fig. 2A2 and B). At 2.5 mM [K^+^]_e_, V_m_ was −73.4 ± 1.37 mV (orange, *N* = 7) and mEPSC frequency normalized to the data at −70 mV was 0.93 ± 0.04 (*N* = 7). Normalized mEPSC frequency at 1 mM [K^+^]_e_ (V_m_ = −85.33 ± 1.9 mV, light orange) was reduced to 0.72 ± 0.03 (*N* = 6), while the frequency at 5 mM [K^+^]_e_ (V_m_ = −54.67 ± 2.72 mV, dark orange) increased to 1.43 ± 0.16 (*N* = 7). The relationship between V_m_ and the mEPSC frequency shown in Fig. 2B indicates that this relationship is not affected by the method of changing V_m_.

**Figure 2.**
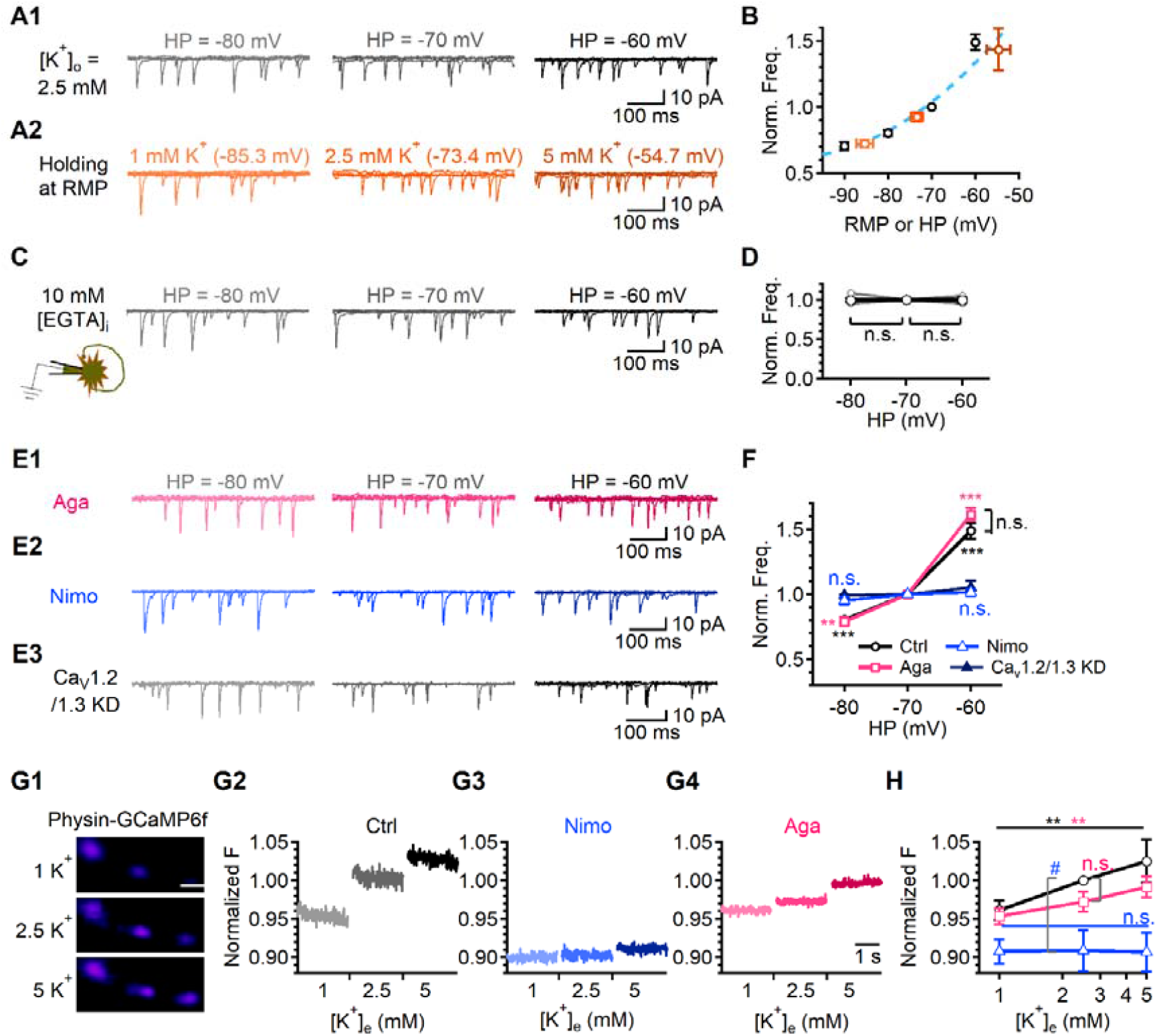
V_m_-dependent regulation of spontaneous release is mediated by LTCCs-dependent changes in basal Ca^2+^. (A). Representative traces of mEPSC frequency in each HP at 2.5 mM [K^+^]_e_ (A1) and each [K^+^]_e_ at the corresponding RMP (A2). (B) A graph indicating the relationship between HP (black) or RMP (orange) and the normalized mini frequency, compared to control −70 mV or 2.5 mM [K^+^]_e_ values. (C) Left. A schematic image of autpase containing 10 mM EGTA patch pipette solution. Right. Representative traces of mEPSC frequency in each HP at 2.5 mM [K^+^]_e_. (D) A graph indicating the average value of the normalized mEPSC frequency in various HP, compared to −70 mV. (E) There are representative traces of mEPSCs in the Aga (E1), Nimo (E2), and shCa_V_1.2/1.3 (E3) in each HP at at 2.5 mM [K^+^]_e_. (F) A graph indicating the average value of the normalized mEPSC frequency in various HP with Aga, Nimo, and shCa_V_1.2/1.3, compared to −70 mV. (G1) Representative resting images of synaptophysin-GCaMP6f (Physin-GCaMP6f) in control condition at different [K^+^]_e_ in the primary cultured hippocampal synapses. Neurons transfected with Physin-GCaMP6f were applied with normal Tyrodes buffer. Scale bar 5 μm. (G2-4) Normalized traces of Physin-GCaMP6f at rest in the various condition of subthreshold potential in control (G2), Nimo (G3), and Aga-treated synapses (G4), compared to control at 2.5 mM [K^+^]_e_. (H) Normalized mean values of fluorescence intensities at resting status in the various condition of subthreshold potential in control, Nimo, and Aga-treated neurons (*Bonferroni* test after *one-way ANOVA*). The individual raw values are described in table S1.

We previously reported that Ca^2+^-dependent spontaneous release in cultured hippocampal neurons is mediated by local Ca^2+^ increase via the stochastic opening of P/Q-, N-, and R-type VGCCs (30). However, 10 mM EGTA in pipette solution completely abolished the effect of subthreshold V_m_ on mEPSC frequency (Fig. 2C and D), suggesting that the V_m_-dependent increase in the spontaneous release is not mediated by nanodomain Ca^2+^ increase induced by the increased open probability of P/Q-, N-, and R-type VGCCs. Accordingly, the V_m_-dependent regulation of spontaneous release was not affected by blocking P/Q-type VGCCs using 100 μ M ω-agatoxin (Aga, Fig. 2E1 and F). Thus, we tested the involvement of global Ca^2+^ changes, possibly attributable to other types of VGCCs such as LTCCs or T-type VGCCs. V_m_-dependent changes in mEPSC frequency were completely abolished by blocking LTCCs using Nimo (Fig. 2E2 and F) or expression of shCa_V_1.2 and shCa_V_1.3 (Fig. 2E3 and F), but not by a low concentration of NiCl_2_ (40 μM), which blocks T-type Ca^2+^ channels (Supplementary Fig. 7). These results demonstrate that LTCCs specifically mediate the V_m_-dependent regulation of spontaneous release. Given that V_m_-dependent regulation of transmitter release by LTCCs was abolished by a high concentration of EGTA, it can be hypothesized that Ca^2+^ influx through LTCCs increases global Ca^2+^ levels at presynaptic terminals and indirectly modulates neurotransmitter release triggered by local Ca^2+^ increases via P/Q-, N-, and R-type VGCCs. To test this hypothesis, we investigated whether presynaptic Ca^2+^ levels are indeed changed by V_m_ and, if this is the case, whether V_m_-dependent changes in Ca^2+^ are affected by LTCC inhibition. To visualize presynaptic Ca^2+^ levels, we expressed GCaMP6f (a genetic Ca^2+^ indicator) fused to synaptophysin (a key synaptic vesicle protein) in hippocampal neuron culture (Physin-GCaMP6f, Fig. 2G1). To estimate changes in basal Ca^2+^ at the resting state ([Ca^2+^]_b_) in presynaptic terminals by V_m_ changes, the fluorescence intensity in the resting condition (F) was measured in different [K^+^]_e_ and normalized to control values obtained in 2.5 mM [K^+^]_e_. F was decreased by hyperpolarization ([K^+^]_e_ = 1 mM) and increased by depolarization ([K^+^]_e_ = 5 mM) under control conditions (black, Fig. 2G2 and H). Nimo reduced F by 9.1 ± 2.97% at 2.5 mM [K^+^]_e_, and V_m_-dependent changes in F were abolished by Nimo (blue, Fig. 2G3 and H). In contrast, Aga did not significantly affect V_m_-dependent changes in F (pink, Fig. 2G4 and H). These results support the hypothesis that V_m_-dependent regulation of presynaptic Ca^2+^ is specifically mediated by LTCCs, which in turn regulate spontaneous transmitter release.

### LTCCs and P/Q-type VGCCs contribute to V_m_-dependent regulation of evoked release with different mechanisms

Next, we examined whether evoked release is also affected by subthreshold V_m_ changes. Measuring the eEPSC amplitude at different V_m_ is not suitable in the autaptic cultured neuron because the eEPSC amplitude of autaptic neurons is affected not only by transmitter release from presynaptic terminals but also by the V_m_ of the postsynaptic compartment. To circumvent this caveat, we used a pHluorin (pH-sensitive GFP)-based assay system that visualizes the amount of vesicle exocytosis by the increase in the fluorescence intensity while V_m_ was changed by changing RMP with different [K^+^]_e_ (Fig. 3A). A single stimulus (1 AP) was applied to primary cultured hippocampal neurons expressing vGlut1-tagged pHluorin (vG-pH) by Ca^2+^-phosphate transfection, and the increase in fluorescence intensity (ΔF) by 1 AP was quantified as a measure of the evoked release amount. The ΔF values obtained under experimental conditions were normalized to the values obtained under normal [K^+^]_e_ conditions (2.5 mM). Hyperpolarization of V_m_ in 1 mM [K^+^]_e_ decreased ΔF by 26.9 ± 6.2% (*N* = 11), whereas depolarization of V_m_ in 5 mM [K^+^]_e_ increased ΔF by 32.8 ± 7.4% (*N* = 11, black, Fig. 3B2 and C) showing the increased evoked release by V_m_ depolarization in control. At 2.5 mM [K^+^]_e_, Aga and Nimo decreased ΔF by 36.3 ± 3.15% and 21.16 ± 5.55%, respectively (Aga, *N* = 11; Nimo, *N* = 10, Fig. 3B1 and C). In the presence of Aga and Nimo, the V_m_-dependent effect on the evoked release was significantly attenuated. A five-fold change in [K^+^]_e_ from 1 to 5 mM induced an increase in ΔF by 1.82-fold in the control but 1.36-fold in Aga (*P* = 0.0464, Fig. 3B2 and C) and 1.33-fold in Nimo (*P* = 0.0225, Fig. 3B2 and C). These findings revealed that both LTCCs and P/Q-type VGCCs contribute to the V_m_-dependent effect on evoked release.

To understand whether two types of VGCCs contribute to V_m_-dependent regulation of evoked release with different mechanisms, we examined effects of [K^+^]_e_ on AP-induced Ca^2+^ increase in presynaptic terminals (peak Ca^2+^) in Physin-GCaMP6f expressing neuron culture (Fig. 3D). A five-fold change in [K^+^]_e_ from 1 to 5 mM induced an increase in Δ[Ca^2+^] by 1.31-fold in the control (Fig. 3E1 and F), which was smaller than the effect of [K^+^]_e_ on glutamate release shown above. Both Nimo and Aga significantly reduced peak Ca^2+^, but the effect of [K^+^]_e_ on peak Ca^2+^ was not affected by Nimo (1.28-fold, Fig. 3E2 and F) but reduced by Aga (1.18-fold, Fig. 3E3 and F), suggesting that LTCCs did not contribute to the increase in AP-driven Ca^2+^ influx by subthreshold depolarization, while P/Q-type VGCCs did, at least in part. These results implied that the Nimo and Aga differentially affect the relationship between AP-induced glutamate release and peak Ca^2+^. To examine this idea, we compared the effect of [K^+^]_e_ on peak Ca^2+^ (solid line) with that on evoked release (broken line) in the control (Fig. 3G1) and in the presence of Nimo (Fig. 3G2), or Aga (Fig. 3G3), in the plots where values are normalized to the values obtained at 2.5 mM [K^+^]_e_ in each experimental condition. The effect of [K^+^]_e_ changes on evoked release was much larger than that on peak Ca^2+^ in control condition (Fig. 3G1). This tendency was maintained in the presence of Aga (Fig. 3G3), but V_m_-dependent effect on release and that on Ca^2+^ were almost identical in the presence of Nimo (Fig. 3G2). Log-log plot between glutamate release and peak Ca^2+^ showed that data points obtained in the control can be fitted by the line with slope of 2.2 (Black dotted line, circle. Fig. 3H), indicating a high Ca^2+^-cooperativity of the release as was reported previously in hippocampal neurons (30, 34). In the presence of Aga, Ca^2+^-cooperativity showed comparable to that of control (1.9, pink dotted line, square, Fig. 3H), suggesting that P/Q-type VGCCs contribute to both AP-evoked Ca^2+^ increase and the release, but do not significantly affect the relationship between two. In the presence of Nimo, however, the slope decreased to 1.1 (blue dotted line, triangle, Fig. 3H), suggesting that LTCC-dependent mechanisms contribute to the high Ca^2+^-cooperativity of the release. Considering the specific role of LTCCs in V_m_-dependent regulation of basal Ca^2+^, it can be suggested that LTCCs-mediated increase in basal Ca^2+^ contribute to the high Ca^2+^-cooperativity of AP-evoked transmitter release.

**Figure 3.**
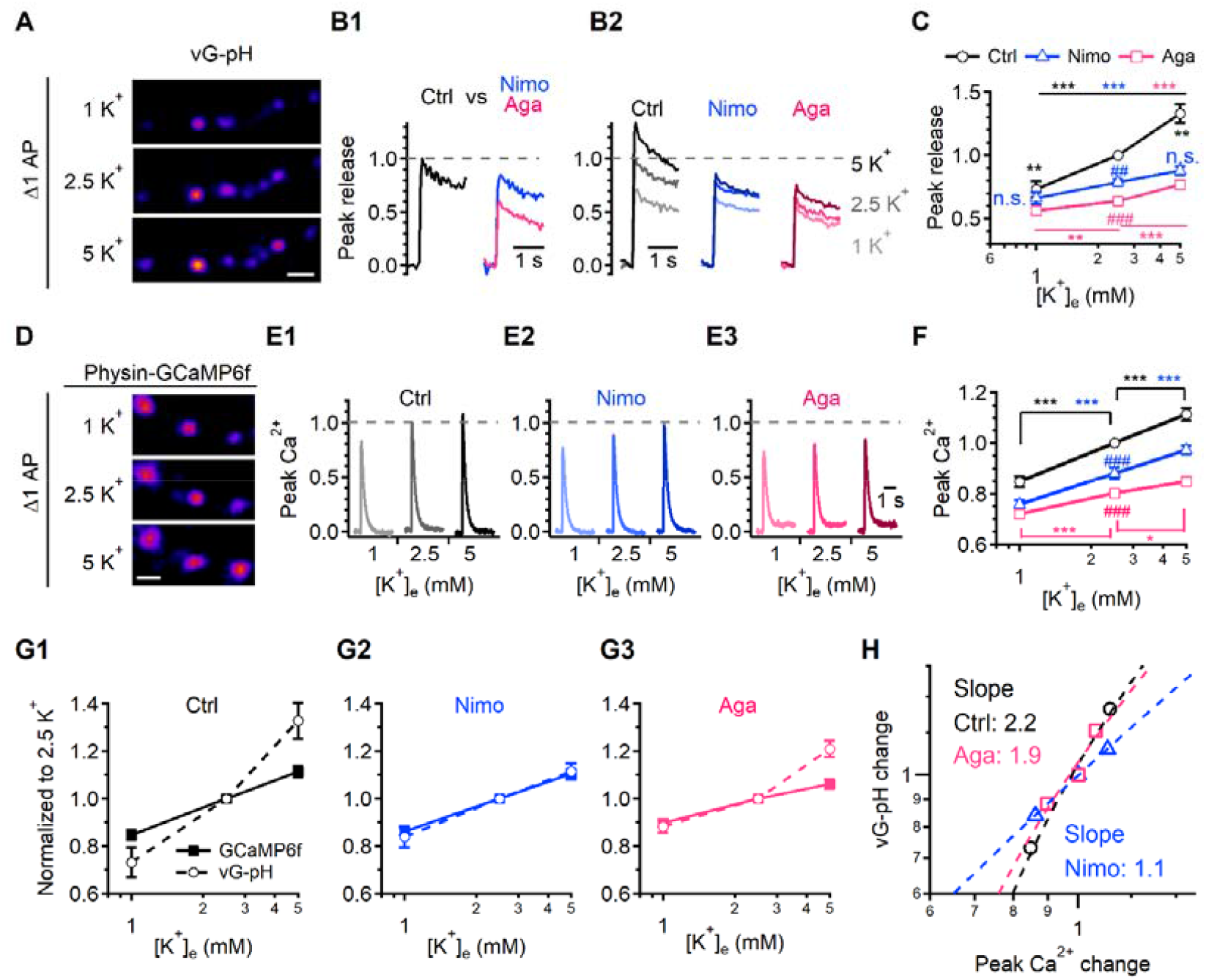
LTCCs and P/Q-type VGCCs contribute to V_m_-dependent regulation of evoked release with different mechanisms. (A) Representative synapse images of vG-pH at 1 AP-evoking (Δ1 AP) neurons in control condition under different [K^+^]_e_. Neurons transfected with vG-pH were stimulated with 1 AP. Scale bar 5 μm. (B1) Ensemble average trances of vG-pH at 1 AP in the control (left, black), Nimo (right, blue), and Aga (right, pink) conditions. (B2) Ensemble average trances of vG-pH at 1 AP at different [K^+^]_e_ conditions of subthreshold potential in control, Nimo, and Aga-treated neurons. (C) Normalized mean values of amplitudes of 1 AP responses in the various subthreshold potential of control, Nimo, or Aga-treated neurons. (D) Representative Δ1 AP images of Physin-GCaMP6f in control condition at different [K^+^]_e_ in the primary cultured hippocampal synapses. (E) Ensemble average trances of Physin-GCaMP6f at 1 AP in the various conditions of subthreshold V_m_ in control (E1), Nimo (E2), and Aga-treated neurons (E3). (F) Normalized mean peak values of amplitudes of 1 AP responses in the various conditions of subthreshold potential in control, Nimo, and Aga-treated neurons. (G) The relationship of [K^+^]_e_ versus the normalized GCaMP6f or vG-pH in control (G1), Nimo (G2), and Aga (G3) condition. (H) A log-log plot for vG-pH change against peak Ca^2+^ change in control (black), Nimo (blue), and Aga (pink) condition. The black dashed line (slope: 2.2), blue dashed line (slope: 1.1), and pink dashed line (slope: 1.9) were fitted by control, Nimo or Aga, respectively. The individual raw values are described in table S1.

### V_m_- and LTCC-dependent regulation of release is mediated by Ca^2+^/calmodulin and PLC-signaling

To explore the signaling mechanism underlying LTCC-mediated regulation of transmitter release, we first investigated the role of Ca^2+^/calmodulin (CaM)-dependent signaling using a calmodulin inhibition peptide (CaM-ip, 10 μM) in a pipette solution. Immediately after patch break-in, mEPSCs (Fig. 4A1 (circle) and A2) and eEPSCs (Fig. 4A1 (triangle) and A3) were recorded sequentially until the effects of CaM-ip perfusion reached a steady state. At the steady state, CaM-ip reduced the mEPSC frequency to 0.77 ± 0.04 (pale bar, Fig. 4B, *N* = 11) and eEPSC amplitude to 0.77 ± 0.07 (solid bar, Fig. 4B, *N* = 10) which was accompanied by the increased PPR (Fig. 4C), while scramble sequence of CaM-ip had no influence (Supplementary Fig. 8), indicating the involvement of Ca^2+^/CaM signaling in transmitter release at physiological RMP. In the presence of CaM-ip, the application of Nimo or Bay K did not show further effects on mEPSC frequency (Fig. 4D and E), eEPSC amplitude (Fig. 4F and G), and PPR (Fig. 4H). Furthermore, V_m_-dependent changes in mEPSC frequency were also abolished (Fig. 4I and J), supporting the idea that LTCC-mediated enhancement of transmitter release is mediated by the increase in [Ca^2+^]_b_.

**Figure 4.**
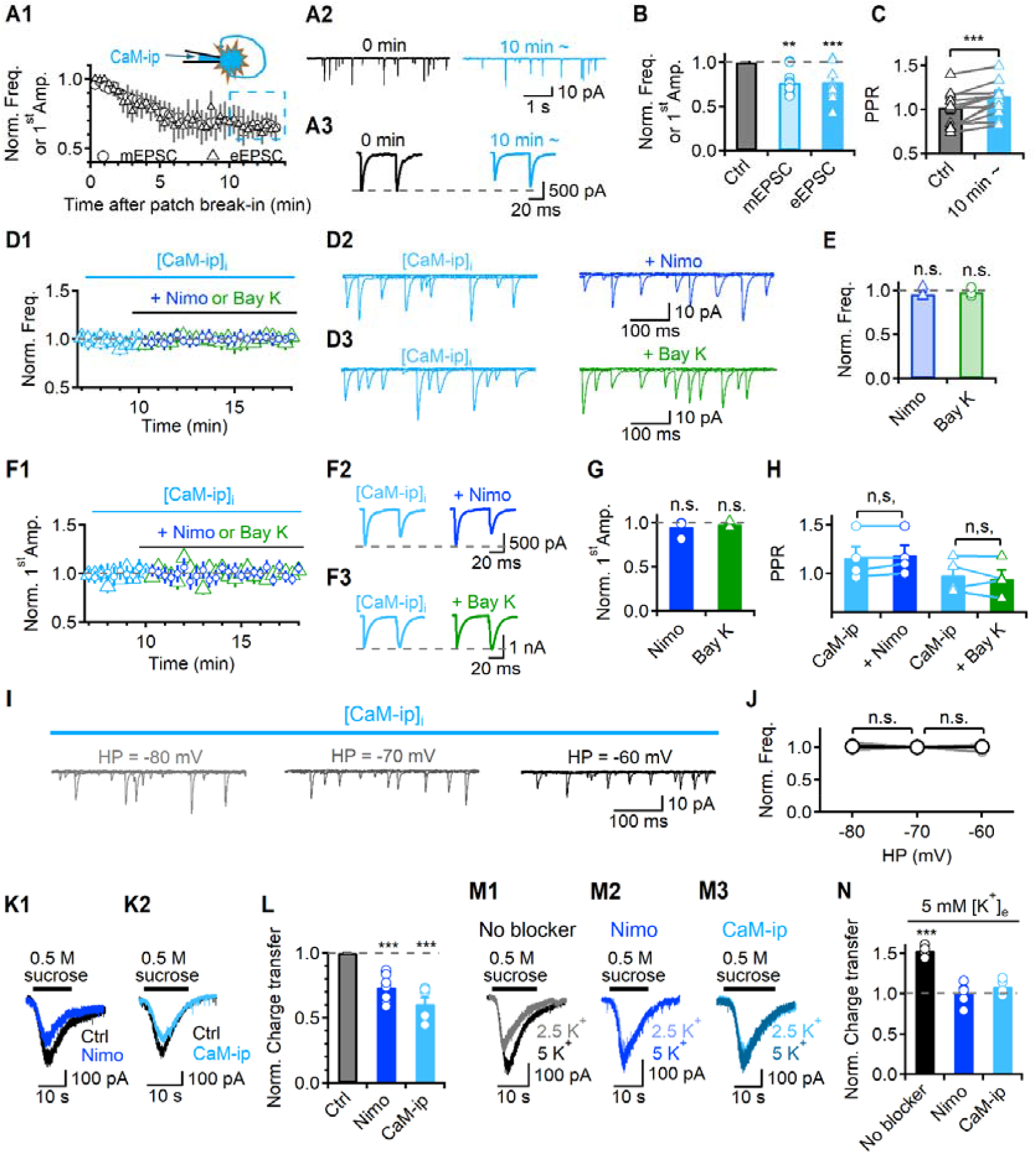
V_m_- and LTCC-dependent regulation of release is mediated by Ca^2+^/calmodulin signaling. (A) A1. Top. A schematic image of CaM-ip diffusion through the autaptic cell body to the axon terminal. Bottom. Average time courses of the normalized mEPSC frequency (circle) or first eEPSC amplitude (triangle) after patch break-in. The data were normalized by the mean frequency or mean amplitude of the first eEPSCs of the initial 1 min after patch break-in, respectively. The light blue dashed line box indicates the steady state average effects, which were used for the bar graphs. (A2, 3) There are representative traces of mEPSCs (A2) and eEPSCs (A3) immediately patch break-in and 10 min after patch in the presence of 10 μM CaM-ip (light blue). The grey dashed line indicates the control first eEPSC peak amplitude. (B) A bar graph showing the normalized mEPSC frequency (pale) and the first eEPSC amplitude (solid) 10 min after patch in the presence of CaM-ip, compared to immediately after patch break-in. A dashed grey line indicates the control level. (C) A bar graph of the average values of the PPR. (D1, F1) An average time course of the normalized mEPSC frequency (D1) and the normalized first eEPSC amplitude (F1) in the presence of CaM-ip followed by Nimo (circle) or Bay K (triangle). (D2-3, F2-3) Representative traces of mEPSC or eEPSC 10 min after patch in the presence of CaM-ip followed by Nimo (D2 and F2) or Bay K (D3 and F3), respectively. (E, G) A bar graph of average value of the normalized mEPSC frequency (E) and the first eEPSC amplitude (G) in the presence of CaM-ip followed by Nimo or Bay K, compared to 10 min after patch break-in. (H) A bar graph of average values of the PPR in different conditions. (I) There are representative traces of mEPSCs in each HP 10 min after patch in the presence of CaM-ip. (J) A graph showing the average value of the normalized mEPSC frequency in various HP in the presence of CaM-ip, compared to −70 mV. (K) Representative traces of the hypertonic sucrose solution application in the presence of each blocker (K1, Nimo; K2, CaM-ip). Top solid line in each trace indicates sucrose application periods. (L) A bar graph of the average values of the normalized charge transfer in different conditions. (M) Representative traces of the hypertonic sucrose solution application for the comparison of 2.5 mM and 5 mM [K^+^]_e_ in the pretreatment of each blocker (M1, no blocker; M2, Nimo; M3, CaM-ip). (N) A bar graph of the average values of the normalized charge transfer in pretreatment of each blocker in 5 mM [K^+^]_e_, compared to 2.5 mM [K^+^]_e_. The individual raw values are described in table S1.

We also tested the role of phospholipase C (PLC) using its inhibitor, U73122. U73122 (2 μM) reduced mEPSC frequency to 0.78 ± 0.04 (Supplementary Fig. 9A1 (circle) and B, *N* = 8) and eEPSCs amplitude to 0.45 ± 0.06 (Supplementary Fig. 9A1 (triangle) and B, *N* = 9) which was accompanied by the increased PPR (Supplementary Fig. 9C), suggesting that PLC-signaling at −70 mV has a significant influence on glutamate release. In the presence of U73122, Bay K showed no further effect on the mEPSC frequency (Supplementary Fig. 9D and E), eEPSC amplitudes (Supplementary Fig. 9D and E), and PPR (Supplementary Fig. 9F). Furthermore, different HP did not alter mEPSC frequency (Supplementary Fig. 9G and H). Taken together, these findings show that Ca^2+^/CaM- and PLC-signaling mediates LTCC-dependent regulation of transmitter release.

The release probability (*p*_r_) and size of the readily releasable synaptic vesicle pool (RRP) are the key determinants of presynaptic neurotransmitter release. Nimo, CaM-ip, and U73122 decreased eEPSC amplitude in association with increased PPR (Fig. 1E-F, 4B-C and Supplementary Fig. 9B-C), suggesting the involvement of increased *p*_r_ in LTCC-mediated facilitation of transmitter release. To investigate whether the V_m_-dependent regulation of transmitter release involves changes in RRP size, we used the hypertonic sucrose solution (500 mM) application technique (35, 36), where the area of the current trace under the baseline during a brief application of sucrose (∼ 15 s) can be regarded as RRP size (Fig. 4K). After patch-break in, the hypertonic sucrose solution was applied with fast-flow rate approximately 1.5 ml min^−1^. Nimo decreased the RRP size by 0.73 ± 0.03 (Fig. 4K1 and L, *N* = 8), suggesting the involvement of RRP in LTCC-mediated regulation of transmitter release. U73122 and CaM-ip in the pipette solution also significantly decreased the RRP size (Fig. 4L and supplementary Fig. 9J, CaM-ip, 0.61 ± 0.05, *N* = 6; U73122, 0.77 ± 0.04, *N* = 8). We then examined whether subthreshold depolarization affected RRP size. At 5 mM [K^+^]_e_, RRP size increased 1.53-fold (*N* = 4, Fig. 4M1 and N), and this increase was not observed in the presence of Nimo, CaM-ip, and U73122 (Fig. 4M, N, and supplementary Fig. 9L). These results suggest that the RRP size is increased by depolarization via Ca^2+^-dependent signaling, which underlies the V_m_-dependent regulation of transmitter release identified in the present study (summarized in supplementary Fig. 10).

We recapitulated the presence and role of LTCCs in the V_m_-dependent regulation of glutamate release in mossy fiber (MF)-CA3 synapses in acute brain slices. GABAergic currents were blocked by picrotoxin. In CA3-PCs, mEPSCs were recorded in the presence of TTX, whereas eEPSCs were recorded by stimulating MF every 20 s with the electrode placed in stratum lucidum. According to the sensitivity to DCG-IV (37) and rise time analysis (38, 39), EPSCs recorded in CA3-PCs were identified to be mainly attributed by MF-CA3 synapses (Supplementary Methods and Supplementary Fig. 11). In the control, Nimo significantly decreased mEPSC frequency (Fig. 5A1 and B, blue, 0.72 ± 0.04, *N* = 15), whereas Bay K increased mEPSC frequency (1.73 ± 0.06, *N* = 4, Supplementary Fig. 12E2). Calmidazolium (CMZ, 10 μM) and U73122 caused a significant decrease in mEPSC frequency (Fig. 5A and B, CMZ, brown, 0.55 ± 0.05, *N* = 12; U73122, orange, 0.78 ± 0.02, *N* = 4). We observed the 1.76-fold increase of mEPSCs frequency by depolarization through increasing [K^+^]_e_ to 5 mM from 2.5 mM (Fig. 5C1 and D, *N* = 7, black). However, this increase was almost completely abolished in the presence of Nimo, CMZ, or U73122 (Fig. 5C and D). The effects of Nimo, Bay K, CMZ, and U73122 on evoked release were similar to those obtained for spontaneous release. Nimo, CMZ, and U73122 significantly decreased eEPSC amplitude (Fig. 5E, F, and I1, Nimo, 0.6 ± 0.04, *N* = 4; CMZ, 0.62 ± 0.07, *N* = 6; U73122, 0.61 ± 0.06, *N* = 9), which was accompanied by the increased PPR (Fig. 5J), whereas Bay K Increased eEPSC amplitude (1.5 ± 0.09, *N* = 5, Supplementary Fig. 12G2) which was accompanied by the decreased PPR (Supplementary Fig. 12G3). The amplitude of eEPSC increased by 1.86-fold in 5 mM [K^+^]_e_, and this increase was almost completely abolished in the presence of Nimo, CMZ, or U73122 (Fig. 5G, H, and I2). To examine the possible involvement of postsynaptic LTCCs in the effects of LTCC targeting drugs on mEPSCs and eEPSCs, we tested the effects of Bay K while postsynaptic LTCCs were inhibited by internal patch pipette solution containing 1 mM verapamil, which was shown to block LTCCs when intracellularly applied (40). We confirmed the blocking effect of intracellular verapamil on the postsynaptic LTCCs (Supplementary Fig. 12A-D). In the presence of verapamil in CA3-PCs, Bay K still enhanced the eEPSC amplitude (Supplementary Fig. 12H2, 1.87 ± 0.17, *N* = 7) and the mEPSC frequency (Supplementary Fig. 12F2, 1.78 ± 0.03, *N* = 5), excluding the possible involvement of postsynaptic LTCCs. Collectively, these results suggest that Ca^2+^/CaM- and PLC-dependent signaling activated by basal Ca^2+^ increase via presynaptic LTCCs contributes to the V_m_-dependent regulation of transmitter release at MF-CA3 synapses in acute hippocampal slices.

**Figure 5.**
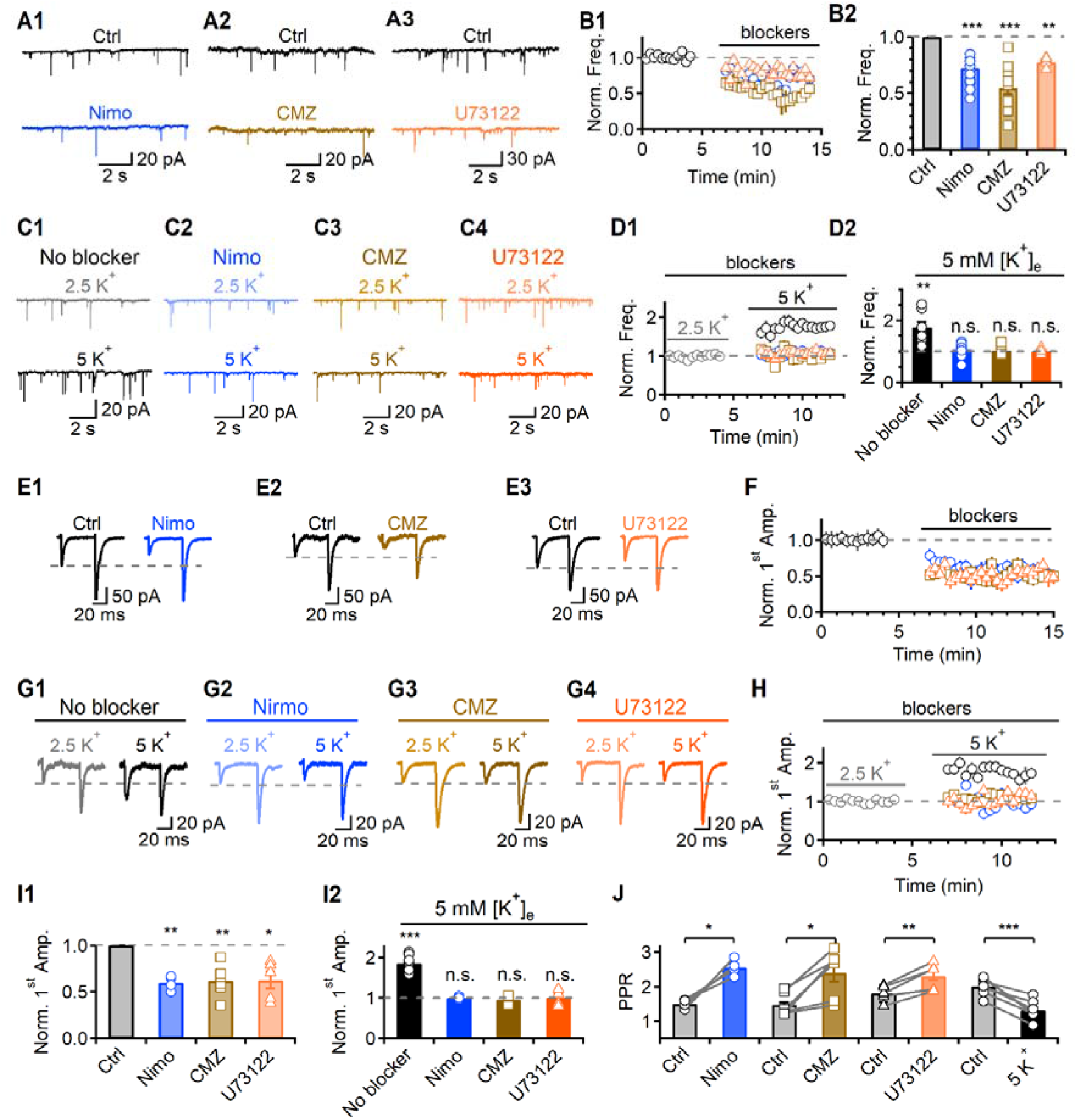
LTCCs mediated V_m_-dependent regulation of glutamate release at the hippocampal MF-CA3 synapses. (A) There are representative traces of mEPSCs in the control, Nimo (A1, blue), CMZ (A2, brown), and U73122 (A3, orange) conditions. (B1) Average time courses of the normalized mEPSC frequency. A dashed grey line indicates the control level. (B2) A bar graph of the average values of the normalized mEPSC frequency in the presence of each blocker in 2.5 mM [K^+^]_e_, compared to 2.5 mM [K^+^]_e_ control. (C) Representative traces of mEPSC for the comparison of 2.5 mM and 5 mM [K^+^]_e_ in the pretreatment of each blocker (C1, no blocker; C2, Nimo; C3, CMZ; C4, U73122). (D1) Average time courses of the normalized mEPSC frequency. (D2) A bar graph of the average values of the normalized mEPSC frequency in pretreatment of each blocker in 5 mM [K^+^]_e_, compared to 2.5 mM [K^+^]_e_. (E) There are representative traces of eEPSCs in the control, Nimo (E1), CMZ (E2), and U73122 (E3) conditions. The grey dashed line indicates the control first eEPSC peak amplitude. (F) Average time courses of the normalized first eEPSC amplitude. (G) Representative traces of eEPSC for the comparison of 2.5 mM and 5 mM [K^+^]_e_ in the pretreatment of each blocker (G1, no blocker; G2, Nimo; G3, CMZ; G4, U73122). (H) Average time courses of the normalized first eEPSC amplitude. (I) A bar graph of the average values of the normalized first eEPSC amplitude in the presence of each blocker in 2.5 mM [K^+^]_e_, compared to 2.5 mM [K^+^]_e_ control (I1) or in pretreatment of each blocker in 5 mM [K^+^]_e_, compared to 2.5 mM [K^+^]_e_ (I2). (J) A bar graph of average values of the PPR in different conditions. The individual raw values are described in table S1.

### Contribution of LTCCs to glutamate release is developmentally regulated

It has been suggested that LTCCs do not participate in neurotransmitter release in most neurons (21, 22). A previous study showed the role of P/Q-, N-, and R-type VGCCs in spontaneous glutamate release at synapses of cultured hippocampal neurons, but the contribution of LTCCs was not recognized (41). Since the neurons that they used were younger than ours (8 to 11 days *vs*. more than three weeks after plating), it is likely that the contribution of LTCC may be developmentally regulated, such that the effect of LTCC did not appear in immature neurons. Therefore, we examined the contribution of each VGCCs, including LTCC, to mEPSCs frequency in hippocampal neurons on days *in vitro* (DIV) 8 to 11 in autaptic cultured neurons. We found that Nimo (Fig. 6A1 and B, *N* = 6) and Bay K (*N* = 4) did not affect mEPSC frequency, whereas Aga, Cono, or 100 μM NiCl_2_ significantly decreased the frequency (Fig. 6A2, supplementary Fig. 13A and B). Consistent with the lack of LTCC contribution, changing the HP between −80 and −60 mV did not affect mEPSC frequency in immature neurons (Fig. 6C, supplementary Fig. 13C, *N* = 6), further supporting the critical role of LTCCs in regulating transmitter release by V_m_.

**Figure 6.**
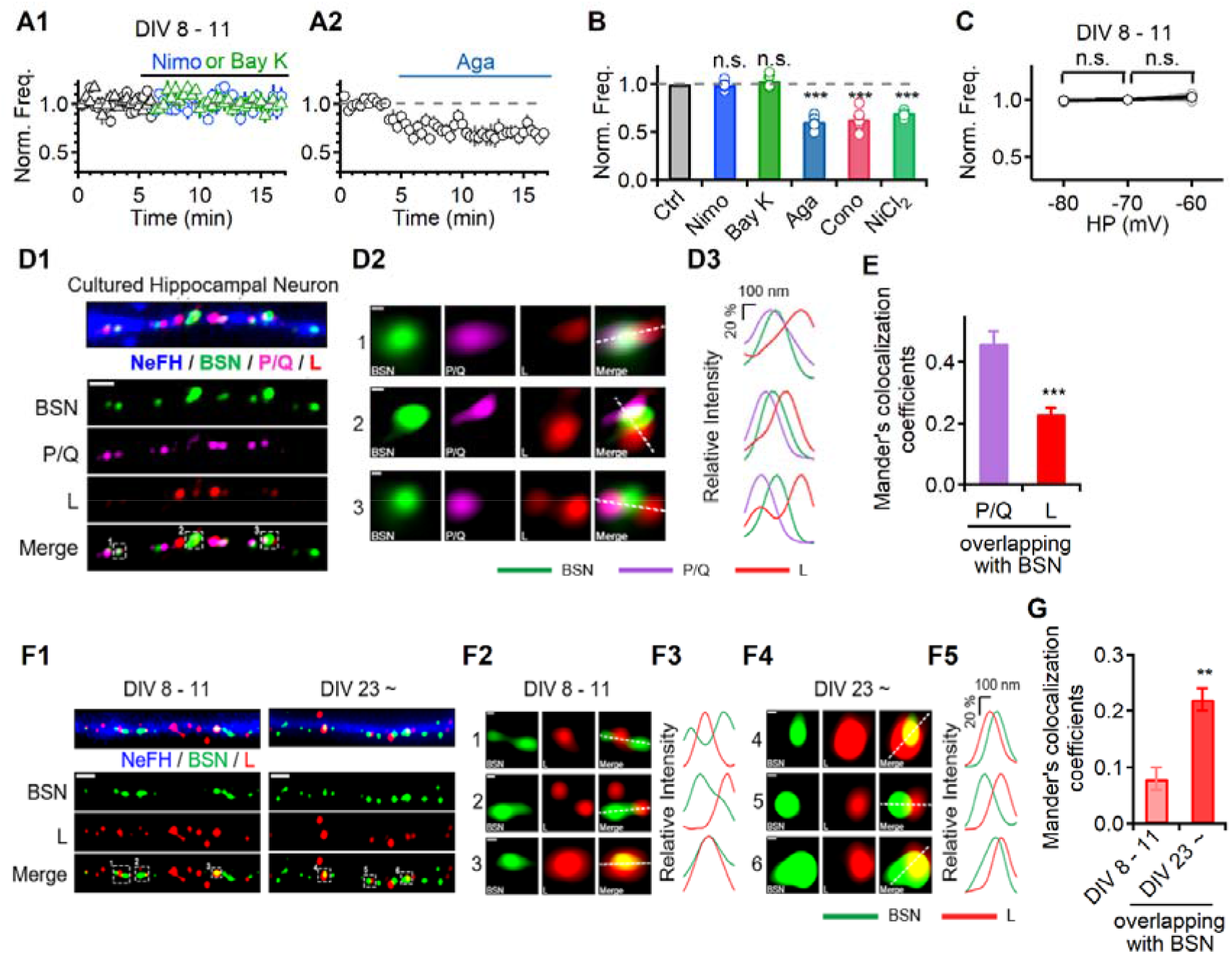
Contribution of LTCCs to glutamate release is developmentally regulated. (A) Average time courses of the normalized mEPSC frequency in different conditions at immature autaptic neurons (A1, blue, Nimo; green, Bay K; A2, Aga). A dashed grey line indicates the control level. (B) A bar graph of average values of the normalized mEPSC frequency in different conditions at immature autaptic neurons, compared to control. (C) A graph indicating the average value of the normalized mEPSC frequency in various HP, compared to −70 mV. (D1) Representative SR images of cultured rat hippocampal neurons at DIV 23 immunostained with NeFH (blue), BSN (green), P/Q-type (magenta), and L-type (red) calcium channels. Scale bar: 1 μm (D2) Magnified axonal bouton images of rectangular areas in D1, Scale bar: 100 nm. (D3) Fluorescent intensity profiles of the respective proteins across the line in axonal boutons in D2. (E) Mander’s colocalization coefficients of each calcium channel overlapped with BSN. (F1) Representative SR images of cultured hippocampal neurons at DIV 8-11 (left) and DIV 23 (right) immunostained with NeFH, BSN, and L-type calcium channel. Scale bar: 1 μm. (F2, F4) Magnified axonal bouton images of rectangular areas in F1, Scale bar: 100 nm. (F3, F5) Fluorescent intensity profiles of the respective proteins across the line in axonal boutons in F2 and F4, respectively. (G) Manders’s colocalization coefficients of the LTCC overlapped with BSN. The individual raw values are described in table S1.

To examine whether the lack of LTCCs contribution to spontaneous release in immature neurons is due to low levels of LTCCs expression in immature neurons, we tested the effect of Nimo on Ca^2+^ currents elicited by depolarization from −70 mV to 0 mV (Supplementary Fig. 14A). A decrease in the peak inward Ca^2+^ current by Nimo observed in immature neurons (Supplementary Fig. 14C and D, 0.39 ± 0.05, *N* = 6) was not significantly different from that in mature neurons (0.35 ± 0.04, *N* = 11, Supplementary Fig. 14D). These results suggest that LTCCs are already present in immature neurons, but their localization to presynaptic terminals may occur later. To examine this idea, we compared the presynaptic expression level of LTCC between immature neurons (DIV 8-11) and mature neurons (DIV 23∼) by analyzing the extent of overlap between LTCCs and the presynaptic marker. We first examined the subcellular localization of LTCC and P/Q-type VGCC along with Neurofilament H (NeFH, an axonal marker) and Bassoon (BSN, an active zone marker) in adult neurons. Airyscan super-resolution (SR) imaging revealed that immunofluorescent signals of P/Q-type VGCC overlapped considerably with those of BSN, whereas the LTCC signals did not fully overlap with those of BSN and most of their localization were in the juxta-synaptic sites (Fig. 6D and E), as previously reported (29). Developmental changes in LTCC density at presynaptic terminals were further confirmed in independent sets of experiments using cultured rat hippocampal neurons and SR imaging showed that the degree of colocalization of LTCC with BSN was significantly higher in adult neurons than in immature neurons (Fig. 6F and G), which is consistent with the results of the electrophysiological data in this study. These results indicate that the presynaptic localization of LTCC is developmentally regulated, which may be why appropriate LTCC-mediated Ca^2+^ signaling to regulate vesicular exocytosis appears in adult neurons.

## DISCUSSION

The molecular mechanisms of neurotransmitter release are well established, but their fine-tuned modulation remains unclear. The present study examined how subthreshold potential changes modulate two classes of synaptic transmission: AP-evoked transmitter release and spontaneous release. First, we found that subthreshold potential changes affected spontaneous as well as AP-evoked release through LTCCs at presynaptic terminals. Second, Ca^2+^ influx through LTCCs did not directly contribute to Ca^2+^-triggered transmitter release, but contributed to the global Ca^2+^ changes in presynaptic terminals that regulate release probability and RRP size via CaM and PLC. Third, we demonstrated that the presynaptic localization and the role of LTCCs in synaptic transmission are developmentally regulated. In the early stages of development, LTCCs hardly co-localized with presynaptic proteins, but their colocalization increased significantly at a more mature stage. Thus, V_m_-dependent regulation of glutamate release requires localization of LTCCs at presynaptic terminals, which appear in the mature state of neurons.

This study raises several interesting questions. The first is how V_m_-dependent changes in presynaptic Ca^2+^ levels in the basal state affect Ca^2+^-triggered transmitter release. Molecular mechanisms mediating Ca^2+^-triggered transmitter release are now well established, in that synaptotagmins bind Ca^2+^ via two C2-domains, and transduce the Ca^2+^ signal into nanomechanical activation of the membrane fusion machinery, the SNARE complex (42). However, it is not well understood how changes in basal Ca^2+^, which are far smaller than AP-evoked local Ca^2+^ increases, regulate transmitter release. Recently, the Ca^2+^-dependence of vesicle priming, fusion, and replenishment was studied in cerebellar mossy fiber boutons, revealing that the number of RRP strongly depends on basal Ca^2+^ between 30 and 180 nM (43). As a potential signaling molecule that regulates Ca^2+^-dependent vesicle priming, the interaction of diacylglycerol/PLC or Ca^2+^/phospholipids with Munc13s (10, 16, 44–46) has been proposed (43). In the present study, we demonstrated that V_m_-induced changes in transmitter release are mediated by basal Ca^2+^ changes and involve CaM and PLC activation (Fig. 2, 4, and supplementary Fig. 9); RRP changes may underlie this effect (Fig. 4 and supplementary Fig. 9). The effect of phorbol esters that potentiate evoked release as well as spontaneous release in the calyx of Held synapses, which was interpreted as an increased fusion “willingness” with the allosteric modulation model (11, 12), may share common mechanisms with V_m_-induced changes shown in the present study.

The role of basal Ca^2+^ changes has been investigated with the aim of explaining synaptic facilitation, leading to the “residual Ca^2+^ hypothesis” (47, 48). Considering that the residual Ca^2+^ induced by high frequency stimulation and the basal Ca^2+^ increase by V_m_ depolarization have different Ca^2+^ sources, namely mitochondrial Ca^2+^ release for the former (49, 50) and LTCCs for the latter (Fig. 2G), it would be intriguing to know whether they share common downstream mechanisms. Mitochondria-dependent residual Ca^2+^ during post-tetanic potentiation was shown to increase the release probability but not RRP size (50). The Ca^2+^-CaM-Munc13-1 complex was suggested to play a pivotal role in short-term synaptic plasticity by regulating the recovery of RRP, but with no changes in RRP size or *p*_r_ at the resting state (9, 10). These differences may imply a difference in downstream mechanisms between V_m_-dependent global Ca^2+^ changes and activity-dependent increases in residual Ca^2+^. However, further studies are required to elucidate the detailed mechanisms. The involvement of distinct Ca^2+^ sensors that detect basal Ca^2+^ changes is of interest. Recently, the involvement of synaptotagmin 7, a high-affinity Ca^2+^ sensor (51), in synaptic facilitation has been suggested (52, 53). Future studies should investigate whether synaptotagmin 7 is also involved in the V_m_-induced changes in transmitter release.

The second question is how presynaptic Ca^2+^ changes that mediate the V_m_-dependent regulation of transmitter release are specifically attributable to LTCCs. L- and T-type VGCCs can open at or near the RMP in CA1 hippocampal neurons (54, 55). According to their biophysical properties, T-type VGCCs, low-voltage activated Ca^2+^ channels, appear to be better suited for contributing to changes in presynaptic Ca^2+^ levels by subthreshold V_m_ changes. However, we found that the blockade of T-type Ca^2+^ channels did not significantly affect the V_m_-dependent regulation of transmitter release (Supplementary Fig. 7). This result suggests that the contribution of VGCCs to the V_m_-dependent regulation of transmitter release does not simply depend on the activation range of VGCCs. Possibly, presynaptic terminals are highly compartmentalized so the localization of the Ca^2+^ source is critical for its function. In fact, the differential roles of L- and T-type Ca^2+^ channels have been demonstrated in the postsynaptic compartment in hippocampal neurons, where L-type, but not T-type Ca^2+^ channels, contribute to metabotropic glutamate receptor 5 (mGluR5)-induced PLC activation, although L- and T-type Ca^2+^ channels equally contribute to depolarization induced Ca^2+^ increase (56). It was thus suggested that LTCCs, but not T-type Ca^2+^ channels, may form a signaling complex with PLC so that LTCC-induced Ca^2+^ microdomains effectively activate PLC (56). The results of the present study, identifying the involvement of PLC (Fig. 5 and supplementary Fig. 9), also suggest a close link between LTCCs and PLC in presynaptic terminals. It is well known that coupling between P/Q-, N-, or R-type VGCCs with low-affinity vesicular Ca^2+^ sensors is critical for Ca^2+^-triggered vesicular fusion (41). Additionally, our study suggests that coupling between LTCCs and unidentified high-affinity Ca^2+^ sensors or Ca^2+^-sensitive signaling proteins such as CaM or PLC may be important for the regulation of release by basal Ca^2+^ changes. Developmental changes in the LTCC contribution to the regulation of transmitter release (Fig. 6A-C) and presynaptic localization (Fig. 6D-G) shown in the present study also highlight the importance of accurate localization for its function.

One of the important points revealed in the present study is that the mechanisms involved in the V_m_-dependent regulation of transmitter release are surprisingly similar between evoked and spontaneous release. Since spontaneous release occurs at basal Ca^2+^ levels in a Ca^2+^-dependent manner, it is generally thought that the Ca^2+^-dependent mechanism underlying spontaneous release is different from that underlying AP-triggered evoked release that occurs in response to a large increase in local Ca^2+^ levels (57). As a candidate for the high-affinity Ca^2+^ sensor responsible for spontaneous release, Doc2 was suggested (58, 59), though the Ca^2+^-dependency of Doc2 is controversial (60). However, several studies have shown that Ca^2+^-dependent component of spontaneous release is not mediated by basal Ca^2+^ but by the local Ca^2+^ increase induced by VGCC openings (30, 41, 61) or Ca^2+^ release from the internal stores (30, 62). If the local Ca^2+^ increase induced by stochastic activation of Ca^2+^ source is sufficient to activate low-affinity Ca^2+^ sensors that mediate AP-evoked transmitter release, spontaneous release may not require high-affinity Ca^2+^ sensors. Consistent with this idea, synaptotagmin-1, a low-affinity Ca^2+^ sensor with high Ca^2+^-cooperativity for fast-evoked release, was shown to mediate spontaneous release in cortical neurons (63). Furthermore, several studies have shown that the apparent Ca^2+^-cooperativity for spontaneous release is not as low as previously thought but similar to that for evoked release (30, 63). Taken together, our study supports the idea that Ca^2+^-dependent mechanisms that operate in the range of presynaptic Ca^2+^ levels in the resting state are not for mediating spontaneous release but for regulating both spontaneous and evoked release. However, this conclusion cannot be generalized to all synapses because Ca^2+^-dependent mechanisms for spontaneous release vary depending on cell type and developmental stage (30). At synapses where spontaneous release is not dependent on VGCCs, such as excitatory synapses in CA1 hippocampal neurons (30) and neocortical neuron cultures (64), spontaneous and evoked release may be regulated by distinct mechanisms.

The RMP is a key component of the physiology of excitable cells, including neurons. The RMP varies widely in different types of neurons and can fluctuate within a subthreshold range under different physiological conditions, including changes in K^+^ concentration, ion channel activity, and synaptic activity. Accumulating evidence demonstrates the importance of neuronal resting Ca^2+^ signaling in the regulation of synaptic efficacy and neuronal homeostasis (65). Basal Ca^2+^-dependent enhancement of transmitter release by subthreshold depolarization has been previously reported in MF-CA3 synapse synapses (2), calyx of Held synapses, and cerebellar interneurons (3). These observations highlight the functional importance of subthreshold potential changes that regulate AP-evoked transmitter release. However, the underlying mechanism of how subthreshold depolarization increases presynaptic Ca^2+,^ which leads to enhancement of transmitter release, was elusive. Furthermore, it was unknown whether subthreshold potential changes influence spontaneous and evoked release via the same mechanism. In the present study, we demonstrated that LTCCs are responsible for the increased presynaptic Ca^2+^ by subthreshold depolarization and that Ca^2+^-dependent signaling, including CaM and PLC, increases RRP size, resulting in the enhancement of both evoked and spontaneous release. Our study highlights the role of LTCC as a key player in the regulation of transmitter release by coupling subthreshold potential changes with the activation of Ca^2+^-dependent signaling molecules that regulate transmitter release.

## METHODS

### Autaptic hippocampal neuron culture and slice preparation

Primary cultures of rat autaptic hippocampal neurons were prepared as described previously with slight adaptations (30, 66). Hippocampal slices were prepared from P20 −30 Sprague-Dawley rats. After anaesthetizing by inhalation with 5% isoflurane, rats were decapitated and the brain was quickly removed and chilled in an ice-cold high-magnesium cutting solution. All preparations were carried out under the animal welfare guideline of Seoul National University (SNU), and approved by IACUC of SNU. The detailed processes are described in supplementary materials and methods.

### Electrophysiology

Autaptic cultured neurons were visualized using an Olympus IX70 inverted microscope and continuously perfused with an extracellular solution. Electrophysiological recordings were performed at room temperature. For recordings from hippocampal slices, slices were transferred to an immersed recording chamber continuously perfused with oxygenated aCSF using a peristaltic pump (Gilson). Temperature was maintained at 35 ± 1 °C. CA3 pyramidal cells were visualized using an upright microscope equipped with differential interference contrast optics (BX51WI; Olympus). Whole-cell voltage-or current-clamp recordings were performed as described in detail in supplementary materials and methods.

### Dissociated hippocampal neuron culture and optical image using vGlut1-pHluorin and synaptophysin-GCaMP6f

Hippocampal CA1-CA3 regions were dissected and dissociated from P0 - P1 SD rats, and plated onto ploy-ornithine-coated glass, as previously described (67). All constructs were transfected 8 days after plating and further incubated for 17 - 25 days in culture media (MEM, 0.5% glucose, 0.01% transferrin, 0.5 mM GlutaMAX-I, 4μM 1-β-D-cytosine-arabinofuranoside, 2% B27, 5% FBS). For presynaptic terminal live imaging to examine synaptic transmission or synaptic Ca2+ levels, vGlut1-pHluorin (vG-pH) or synaptophysin-GcaMP6f (Physin-GcaMP6f) constructs were utilized. For details, please see supplementary materials and methods. The detailed experimental procedures were described in supplementary materials and methods.

### Immunocytochemistry and antibodies

For immunochemistry, dissociated cultured hippocampal neurons were fixed in 4% paraformaldehyde (PFA) in 4% sucrose-containing PBS for 15 min and permeabilized for 5 min in 0.25% Triton X-100 at room temperature. Anti-chicken Neurofilament H (TA309177, Origine), anti-rabbit Bassoon (141 002; Synaptic Systems), anti-guinea pig Cav2.1 (152 205; Synaptic Systems), anti-mouse Cav1.2 (MA5-27717, Invitrogen) were used in the experiments and Alexa Fluor secondary antibodies were purchased from Thermo Fisher Scientific. The detailed processes are described in supplementary materials and methods.

### Drugs

ω-Agatoxin-IVA, ω-Conotoxin GVIA, TTX, and were purchased from Alomone Labs (Jerusalem, Israel). Bay K8644, U73122, and calmidazolium were purchased from Tocris (Bristol, UK). Calmodulin inhibitory peptide and calmodulin inhibitory peptide scramble were purchased from Calbiochem (Darmstadt, Germany). All other chemicals were purchased from Sigma (St. Louis, MO, USA). Toxin stock solutions were made at 1000-fold concentration with distilled water or DMSO and stored at −20°C.

### Statistical analysis

Data were expressed as the mean ± SEM, where *N* represents the number of cells studied. Statistical analysis was performed using IgorPro (version 6.1, WaveMetrics, Lake Oswego, OR, USA) and OriginPro (version 9.0, OriginLab Corp., Northampton, MA, USA). Significant differences between the experimental groups were analyzed using independent or paired Student’s *t*-tests. All data are represented as the mean ± S.E.M., \**P*<0.05, \*\**P*<0.01, \*\*\**P*<0.001; n.s. = not significant.

## Acknowledgments

This research was supported by the National Research Foundation grants from the Korean Ministry of Science and ICT (2021R1A6A3A01088217 to Lee BJ, 2020R1A2B5B02002070 to W.-K. Ho, and 2020R1A2C2010791 to Kim SH). Portions of this paper were developed from the thesis of B. J. Lee.

