## Supplementary information for "L-type Ca^2+^ channels mediate regulation of glutamate release by subthreshold potential changes"

[1]Department of Physiology and Biomedical Sciences, [2]Neuroscience Research Institute, Seoul National University College of Medicine, [3]Interdisciplinary Program in Neuroscience, [4]Department of Brain and Cognitive Science, Seoul National University College of Natural Science, [5]Department of Neuroscience, Graduate School, 6Department of Physiology, School of Medicine, Kyung Hee University, Seoul, Korea

[a]Present address: HHMI Janelia Research Campus,19700 Helix Dr, Ashburn, VA 20147, USA

[b]Present address: Samsung Advanced Institute of Technology, Samsung Electronics, Suwon-si, South Korea

*Corresponding authors

Won-Kyung Ho, M.D., Ph.D.

Department of Physiology, Seoul National University College of Medicine, 103 Daehak-ro, Jongno-gu, Seoul 03080, Korea

Sung Hyun Kim, Ph.D.

Department of Physiology, School of Medicine, Kyung Hee University,

Medical Building Rm 901, 26 Kyungheedae-ro, Dongdaemun-gu, Seoul,02447, Korea

**This PDF file includes:**

Supplementary materials and methods

Figures S1 to S15

Tables S1

SI References

### Supplementary Materials and Methods

#### *Autaptic hippocampal neuron culture and double knockdown of $Ca_V1.2$ and $Ca_V1.3$*

For primary cultures of rat autaptic hippocampal neurons, hippocampal neurons and astrocytes were obtained from Sprague-Dawley (SD) rats according to the protocols approved by the Seoul National University Institutional Animal Care and Use Committee (SNU-210416-2-1). Astrocyte cultures were prepared from the SD rat cortices P0 - P1 and grown for 10 days in 100-mm culture dish in glial medium [minimum essential medium (MEM; Invitrogen) supplemented with 0.6% glucose, 1 mM pyruvate, 2 mM GlutaMAX-I (Invitrogen), 10% horse serum (HS; Invitrogen), and 1% penicillin-streptomycin (PS; Invitrogen)] before plating on the poly-D-lysine (PDL, Sigma, St. Louis, MO, USA) sprayed microisland coverslips in 30-mm petri dishes. 2 - 3 days before neurons being added in sprayed microisland dishes, astrocytes were removed from the 100-mm culture dish using trypsin-EDTA (Invitrogen) and plated on the microisland coverslips at a density of 60,000 cells/dish. Hippocampi from P0 - P1 SD rats were dissected in Hank's balanced salt solution (HBSS, Invitrogen), digested with papain (Worthington, Freehold, NJ, USA), and then triturated with a polished half-bore Pasteur pipette. Immediately after removing glia medium in 30-mm dishes of microisland-shaped astrocytes, hippocampal neurons were added at a density of 6,000 cells/dish and were grown in neurobasal medium supplemented with 2% B27 (Invitrogen) and 0.5 mM GlutaMAX-I.

To knockdown LTCCs, the shRNAs of $Ca_V1.2$ (3) and $Ca_V1.3$ (4) were used as previously reported. The $Ca_V1.2$-targeting shRNA sequence is 5′-GCCGAAATTACTTCAATATTTCAAGAGAATATTGAAGTAATTTCGGC-3′. The $Ca_V1.3$-targeting shRNA sequence is 5′-TTATCTCTCATGGCAACTTTCCCACTTCCTGTCATGGGAAAGTTGCCATGAGAGATAAA-3′. The $Ca_V1.2$ targeting shRNA sequences have been validated in HEK293 cells and primary hippocampal neurons which has been reported previously (3). The $Ca_V1.3$ targeting shRNA sequences have been validated in primary neuronal culture (4). Customized adeno-associated virus (AAV) containing $Ca_V1.2$ shRNA-mCherry and $Ca_V1.3$ shRNA-GFP were produced from the virus facility for brain research institute in Korea institute of science and technology (KIST). A volume of 1 μl (4.0 X 10^12 genome copies/ml) of purified AAV was used to infect autaptic cultured neurons at DIV 15 and experiments were performed at least 15 days after infection. The images of AAV infected autapse were acquired by Olympus inverted fluorescence microscope (IX53, Olympus, Tokyo, Japan) equipped with a 40X objective lens (Olympus, LUCPlanFLN) using Olympus digital camera (Olympus, DP73) driven by CellSens (Olympus) imaging software.

#### *EPSC recordings from autaptic cultured hippocampal neurons*

Autaptic cultured neurons were perfused with an extracellular solution consisting of the following composition (in mM): 135 NaCl, 2.5 KCl, 2 $CaCl_2$, 1 $MgCl_2$, 10 glucose, 10 HEPES, 0.1 EGTA, pH 7.3, adjusted with NaOH (295-300 mOsm), and the rate of perfusion was maintained at 0.5 mL $min^{-1}$. In the hypertonic sucrose application experiments, a fast flow rate (1.5 mL $min^{-1}$) was required. NaCl was reduced to maintain the osmolarity for high $[K^+]_e$ experiments, as needed. All recordings were performed at least 23 days after neurons were plated on coverslips. For the immature neuron experiments, recordings of mEPSCs and $I_{Ca}$ were performed from DIV 8 to 11 neurons, as needed. The internal pipette solution for recording eEPSCs, mEPSCs and RMP was a K-gluconate-based solution containing 130 K-gluconate, 8 NaCl, 4 MgATP, 0.3 $Na_2GTP$, 10 HEPES, 0.1 EGTA, pH 7.3, adjusted with KOH (295-300 mOsm). The unit of the mEPSC frequency is Hertz.

Autaptic eEPSCs were evoked by a 2 ms voltage-clamp step to 0 mV from an HP of -70 mV with pairs of stimuli (at 50 ms interval). Postsynaptic currents were induced after large and brief activation of the $Na^+$ current. The cells were identified as glutamatergic neurons based on the fast decay kinetics of synaptic currents (5). To examine whether the measurement of eEPSC amplitude was affected by voltage clamp error, we followed the protocol described in previous studies (6). The reduction of eEPSC amplitude by low dose of CNQX (0.5 μM) in 1 mM $[Ca^{2+}]_e$ (Supplementary Fig. 15, 0.71 ± 0.03, $N$ = 6) and that in 10 mM $[Ca^{2+}]_e$ (0.69 ± 0.03, $N$ = 6) were not significantly different ($p$ = 0.56), confirming that voltage clamp error was negligible. After 5 min of stabilization from patch break-in, autaptic eEPSCs were recorded every 20 s for 5 min baseline recordings, followed by individual drug application.

The mEPSCs of hippocampal autapses were recorded at an HP of -70 mV. Events exceeding 6 - 7 pA within a specified interval of three-four digitized points (0.5 - 0.8 ms) that showed a single exponential decay time course were identified as mEPSC. The rise time of mEPSC indicated a 20 - 80% rise time. The mEPSC frequency was measured within 20 s bins. From the continuous recordings at -70 mV without stimulation, toxins and chemicals were typically applied for 5 - 20 min until a constant effect was observed. To rule out any effect of $Ca^{2+}$ depletion of internal $Ca^{2+}$ stores on spontaneous release, recordings were completed within 20 min after break-in.

***Hippocampal Slice Preparation and EPSC recordings from MF-CA3 synapses***

The cutting solution for preparing hippocampal slices contained the following composition (in mM): 110 choline chloride, 25 $NaHCO_3$, 20 glucose, 2.5 KCl, 1.25 $NaH_2PO_4$, 1 sodium pyruvate, 0.5 $CaCl_2$, 7 $MgCl_2$, 0.57 ascorbate (pH 7.3 bubbled with 95% $O_2$ - 5% $CO_2$; osmolarity ~300 mOsm. The isolated brain was glued onto the stage of a vibrating blade microtome (VT1200S, Leica Microsystems) and 300 μm-thick transverse hippocampal slices were cut. The slices were incubated at 34 °C for 30 min in artificial cerebrospinal fluid (aCSF) containing the following (in mM): 125 NaCl, 25 $NaHCO_3$, 20 glucose, 2.5 KCl, 1.25 $NaH_2PO_4$, 1 sodium pyruvate, 2 $CaCl_2$, 1 $MgCl_2$, 0.57 ascorbate, bubbled with 95% $O_2$ - 5% $CO_2$, and thereafter maintained at room temperature until required. For recording from hippocampal slices, slices were transferred to an immersed recording chamber continuously perfused with oxygenated aCSF using a peristaltic pump (Gilson). The rate of aCSF perfusion was maintained at 1-1.5 mL $min^{-1}$. The internal pipette solution for recording EPSCs in CA3-PCs was same as the solution used in the autapse recording.

To record eEPSCs of MF-CA3 synapses, we placed a monopolar stimulation electrode filled with a recording solution in glass pipette (1 - 2 MΩ) in stratum lucidum of the hippocampal CA3 region and applied low-intensity stimulation (100 μs duration; 6 - 20 V intensity), while GABAergic synaptic input was blocked using 100 μM picrotoxin. Stimulation pulses were generated using a digital stimulator (WPI DS8000; World Precision Instruments, Sarasota, FL, USA) and fed into the stimulation electrode via an isolation unit (WPI stimulus isolator DLS100). After 5 min of stabilization from patch break-in, MF-CA3 eEPSCs were recorded every 20 s for 5 min baseline recordings, followed by individual drug application. When mEPSCs were recorded, we added 0.5 μM TTX.

To verify whether EPSCs recorded from CA3-PCs are originating from MF, we examined the effect of DCG-IV (7) and EPSC rise time (8, 9). Bath application of DCG-IV significantly decreased the mEPSC frequency by 0.4-fold and there was no further reduction detected with additional application of P/Q, N, and R-type $Ca^{2+}$ channel blockers (Supplementary Fig. 11A), suggesting that most of VGCC-dependent spontaneous glutamate release is occurring selectively at MF terminal-CA3 synapses. Nevertheless, we cannot entirely exclude presence of mEPSCs coming from other synapses, though the proportion is negligible. The rise time of eEPSC ranged from 0.45 to 0.89 ms (0.65 ± 0.02 ms, $N$ = 34, Supplementary Fig. 11C), which is consistent of previous data of ~ 1 ms for EPSCs of MF-CA3 (0.64 -

0.84) (8). The eEPSC amplitudes were reduced by at least 80% by DCG-IV (Supplemental Fig. 11B), supporting that MF was selectively stimulated.

For the high $[K^+]_e$ experiments, NaCl was reduced to maintain the osmolarity. Recordings were made in somata using an EPC-10 amplifier (HEKA Electonik; Lambrecht/Pfalz, Germany). The signals were low-pass filtered at 5 kHz (low-pass Bessel filter) and sampled at 10 kHz. Series resistance ($R_s$) was monitored, and only recordings with $R_s$ remaining constant (<30% change during a recording) were used. $R_s$ was compensated for 50 - 70%. The data were analyzed using IGOR software (Wavemetrics, Lake Oswego, OR, USA). Patch electrodes were pulled from borosilicate glass capillaries to a resistance of 3 - 4 MΩ when filled with the pipette solution.

#### *Optical image using vGlut1-pHluorin and synaptophysin-GCaMP6f*

For presynaptic terminal live imaging to examine synaptic transmission or synaptic $Ca^{2+}$ levels, vGlut1-pHluorin (vG-pH) or synaptophysin-GCaMP6f (Physin-GCaMP6f) constructs were transfected eight days after dissociated hippocampal neuron culture. Experiments were performed at DIV 17 - 25 after plating. The coverslips were mounted in a stimulation chamber with laminar flow perfusion on the stage of a custom-built laser-illuminated epifluorescence microscope. Live images were acquired using an Andor iXon Ultra 897 (Model #DU-897U-CS0-#BV) back-illuminated EM CCD camera. A diode-pumped OBIS 488 laser (Coherent) shuttering by synchronization with the EMCCD camera during acquisition was utilized as the light source. Fluorescence excitation/emission and collection were achieved using a 40x (1.3 NA) Fluar Zeiss objective lens with 500 - 550 nm emission and 498 nm dichroic filters (Chroma) for pHluorin and GCaMP6f. APs were evoked by passing a 1 ms current pulse through platinum-iridium electrodes from an isolated current stimulator (World Precision Instruments). The neurons were perfused with various Tyrode's buffers containing 1, 2.5, or 5 mM KCl. All experiments were performed at 30 °C.

All images were analyzed using Image J (http://rsb.info.nih.gov/ij) with plugin Time Series Analyzer which is available at https://imagej.nih.gov/ij/plugins/time-series.html. All responsive synaptic boutons were selected for analysis. Fluorescence traces were analyzed using Origin Pro 2020. Peak amplitude of each 1 AP (ΔF values of 1 AP response) was normalized using the peak value of 1 AP at $[K^+]_e$ 2.5 mM response.

#### *Immunocytochemistry*

After fixation of dissociated cultured neurons, they were blocked for 1 hr with blocking solution (3% bovine serum albumin (BSA) and 0.05% Triton X-100 in PBS) at room temperature and incubated with respective primary antibodies diluted in blocking solution overnight at 4 °C, followed by incubation with respective secondary antibody in blocking solution for 45 min at 37 °C. After immunostaining, neurons on coverslips were mounted on a glass slide and super-resolution images were acquired by ZEISS laser scanning microscope LSM980 with Airyscan2 (Carl Zeiss, Jena, Germany) equipped with a 63X oil-immersion objective lens (Plan-Apochromat 63x/1.4) using AxioCam305 mono camera (Carl Zeiss) driven by Zen blue 3.4 software. To estimate the protein colocalization, Mander's colocalization coefficients were analyzed by using FIJI/ImageJ software with the JACoP plug-in function. The average of Mander's colocalization coefficients value of each sample was compared by Student's *t*-test, and its values are indicated as mean ± SEM.

### Supplementary Figures and Figure legends

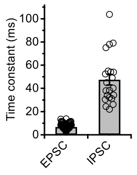


**Supplementary Fig. 1**. The decay time constant of evoked EPSC and IPSC.

A bar graph indicating the decay time constants of evoked EPSC (6.9 ± 0.2 ms, *N* = 94) and IPSC (47.4 ± 4.6 ms, *N* = 21).

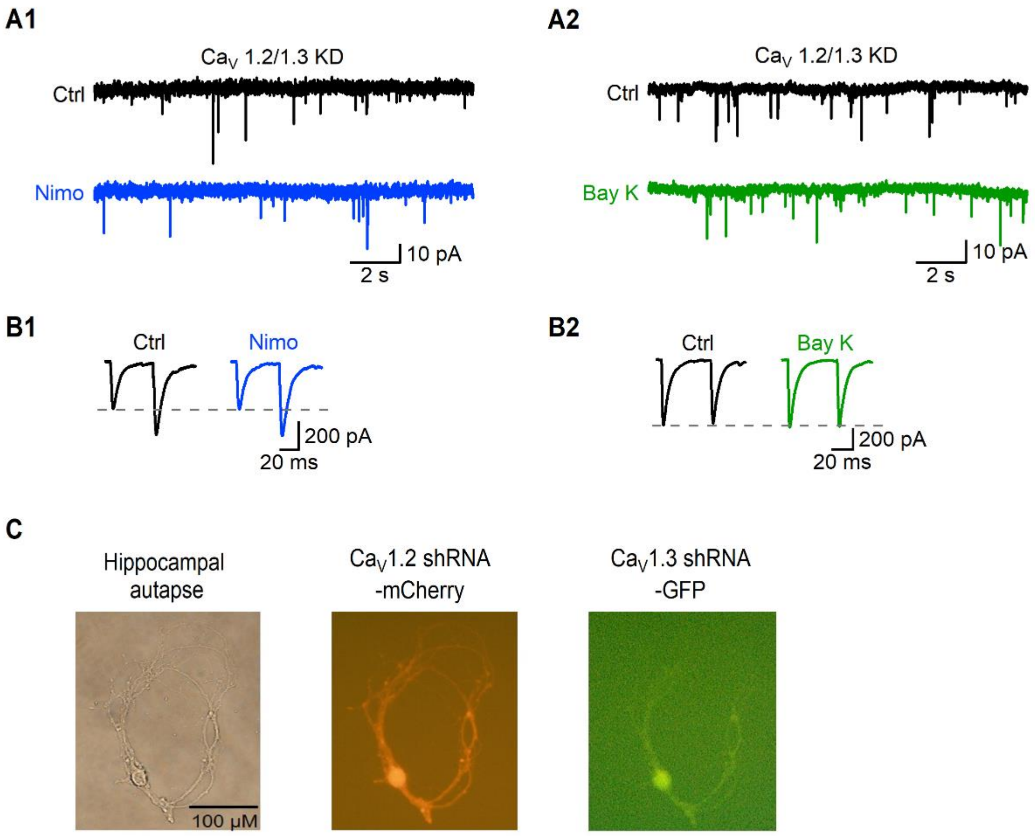


**Supplementary Fig. 2.** Representative traces for figure 1G to 1I.

(A) Representative traces of mEPSCs in the control, Nimo (A1), and Bay K (A2) condition at shCa$_V$ 1.2/1.3 autapses. (B) Representative traces of eEPSCs in the control, Nimo (B1), and Bay K (B2) condition at shCa$_V$ 1.2/1.3 autapses. The grey dashed line indicates the control first eEPSC peak amplitude. (C) Example images of AAV-infected hippocampal autaptic neuron labeled for shCa$_V$1.2

(mCherry, middle), shCa$_V$1.3 (GFP, right), and an identical autapse without fluorescence (Left).

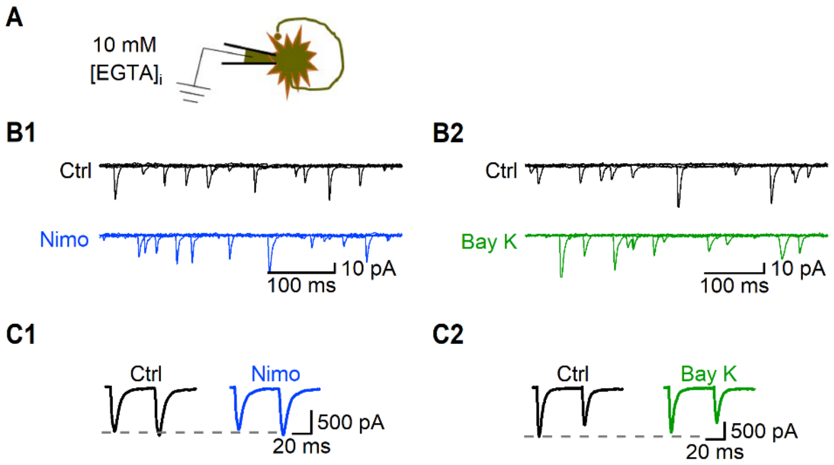


**Supplementary Fig. 3.** Representative traces for figure 1J to 1K.

(A) A schematic image of autaptic pyramidal neuron containing 10 mM EGTA internal patch pipette solution. (B) Representative traces of mEPSCs in the control condition and in the presence of Nimo (B1) or Bay K (B2). (C) Representative traces of eEPSCs in the control condition and in the presence of Nimo (C1) or Bay K (C2). The grey dashed line indicates the control first eEPSC peak amplitude.

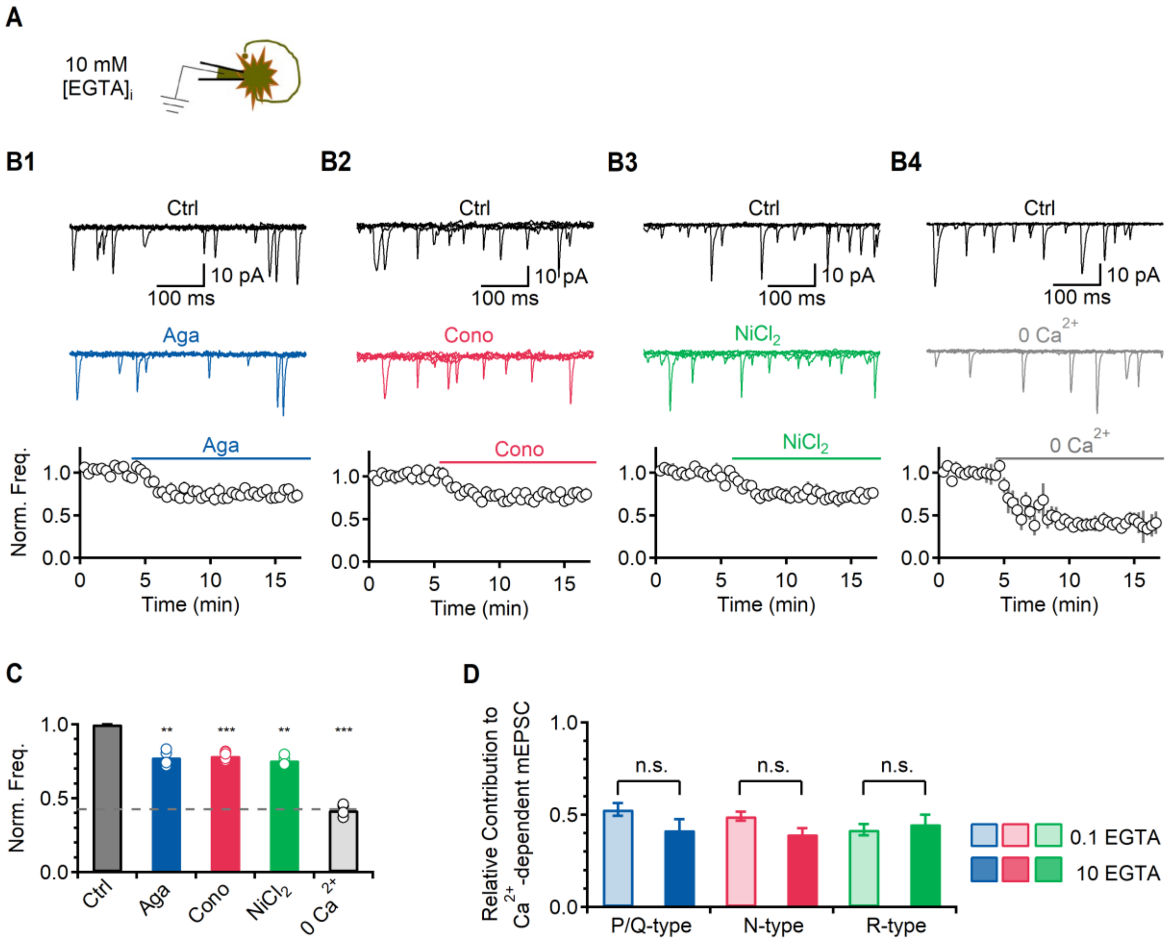


**Supplementary Fig. 4**. The effects of VGCCs or $Ca^{2+}$ removal on the spontaneous release in 10 mM EGTA internal patch pipette solution.

(A) A schematic image of autaptic pyramidal neuron containing 10 mM EGTA internal patch pipette solution. (B) Top. Representative traces of mEPSCs in control, Aga (B1, blue), Cono (B2, red), $NiCl_2$ (B3, green) and removal of $[Ca^{2+}]_e$ (B4). Five 500 ms-long mEPSC traces were overlaid. Bottom. Average time courses of the normalized mEPSC frequency. In each time course plot, the solid lines indicate the presence of each drug or removal of $[Ca^{2+}]_e$. The data were normalized by the mean mEPSC frequency of control. (C) A bar graph of average values of the normalized mEPSC frequency in different conditions, respectively. A dashed line indicates the mEPSC frequency at 0 mM $[Ca^{2+}]_e$. (D) A chart for the VGCC contribution to $[Ca^{2+}]_e$-dependent mEPSCs in 0.1 mM $[EGTA]_i$ (pale bar) or 10 mM $[EGTA]_i$ (solid bar). The individual raw values are described in table S1.

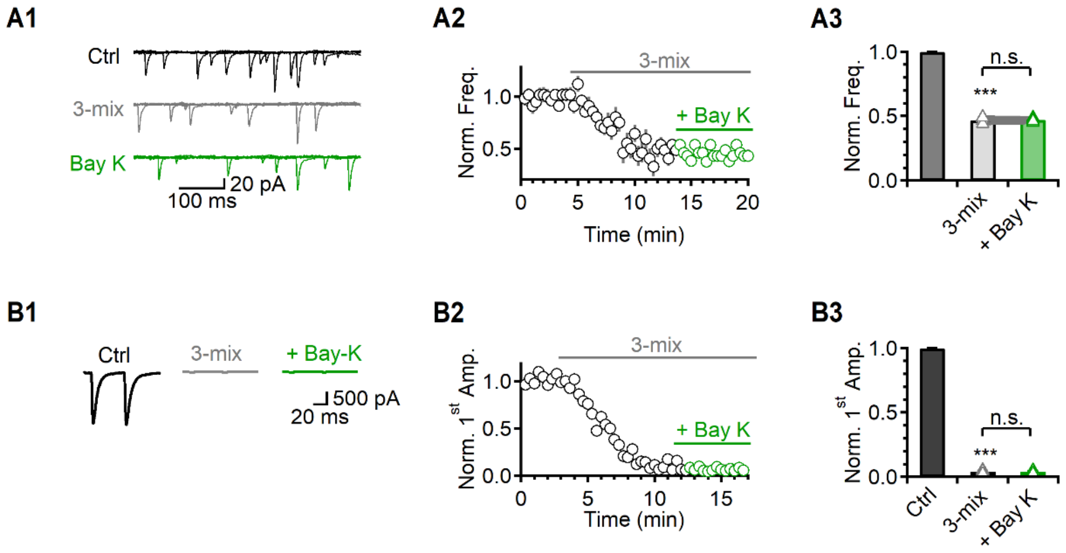


**Supplementary Fig. 5.** The effect of Bay K on spontaneous and evoked glutamate release under blockade of P/Q-, N-, and R-type channels.

(A1, B1) Representative traces of mEPSCs (A1) and eEPSCs (B1) in the control condition and in the presence of 3-mix (gray) followed by Bay K. (A2, B2) Average time courses of the normalized mEPSC frequency (A2) and first eEPSC amplitude (B2). The upper solid line indicates the presence of 3-mix and the lower solid line indicates the additional application of Bay K. (A3, B3) A bar graph of average values of the mEPSC frequency (A3) and first eEPSC amplitude (B3) in presence of 3-mix and additional treatment of Bay K. The individual raw values are described in table S1.

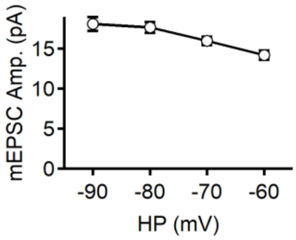


**Supplementary Fig. 6.** The mEPSC amplitude change by $V_m$ change.

A bar graph of the average values of the mEPSC amplitudes with $V_m$ change. The individual raw values are described in table S1.

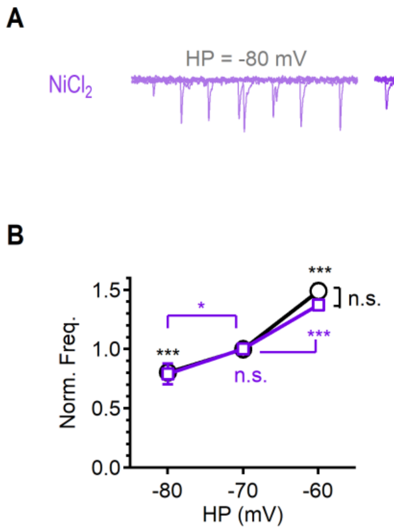


**Supplementary Fig. 7.** The effect of T-type $Ca^{2+}$ channels on mEPSC in various HP.

(A) Representative traces of mEPSC in the presence of 40 μM $NiCl_2$ in each HP at 2.5 mM $[K^+]_e$. (B) A graph indicating the average value of the normalized mEPSC frequency in various HP, compared to control -70 mV. The individual raw values are described in table S1.

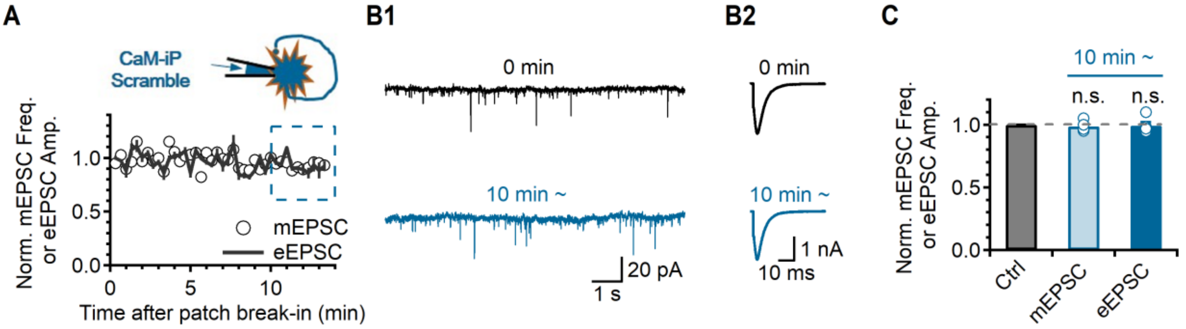


**Supplementary Fig. 8.** The effect of CaM-ip scramble on spontaneous release or evoked release.

(A) Top. A schematic image of CaM-ip scramble diffusion through the autaptic cell body to the axon terminal. Bottom. Average time courses of the normalized mEPSC frequency and the first eEPSC amplitudes after patch break-in. The data were normalized by the mean frequency (circles) of the initial 1 min after patch break-in and the mean amplitude (solid line) of the first 3 eEPSCs after patch break-in. The light blue dashed line box indicates the steady state of drug effects, which were used for the bar graphs. (B) Representative traces for the comparison of mEPSCs (B1) and eEPSCs (B2) between immediately patch break-in (top) and 10 min after (bottom) patch in presence of 10 μM CaM-ip scramble. (C) A bar graph showing the average values of the normalized mEPSC frequency (pale) and first eEPSC amplitudes (solid) 10 min after patch in presence of CaM-ip scramble, compared to immediately after patch break-in. A dashed grey line indicates the control level. The individual raw values are described in table S1.

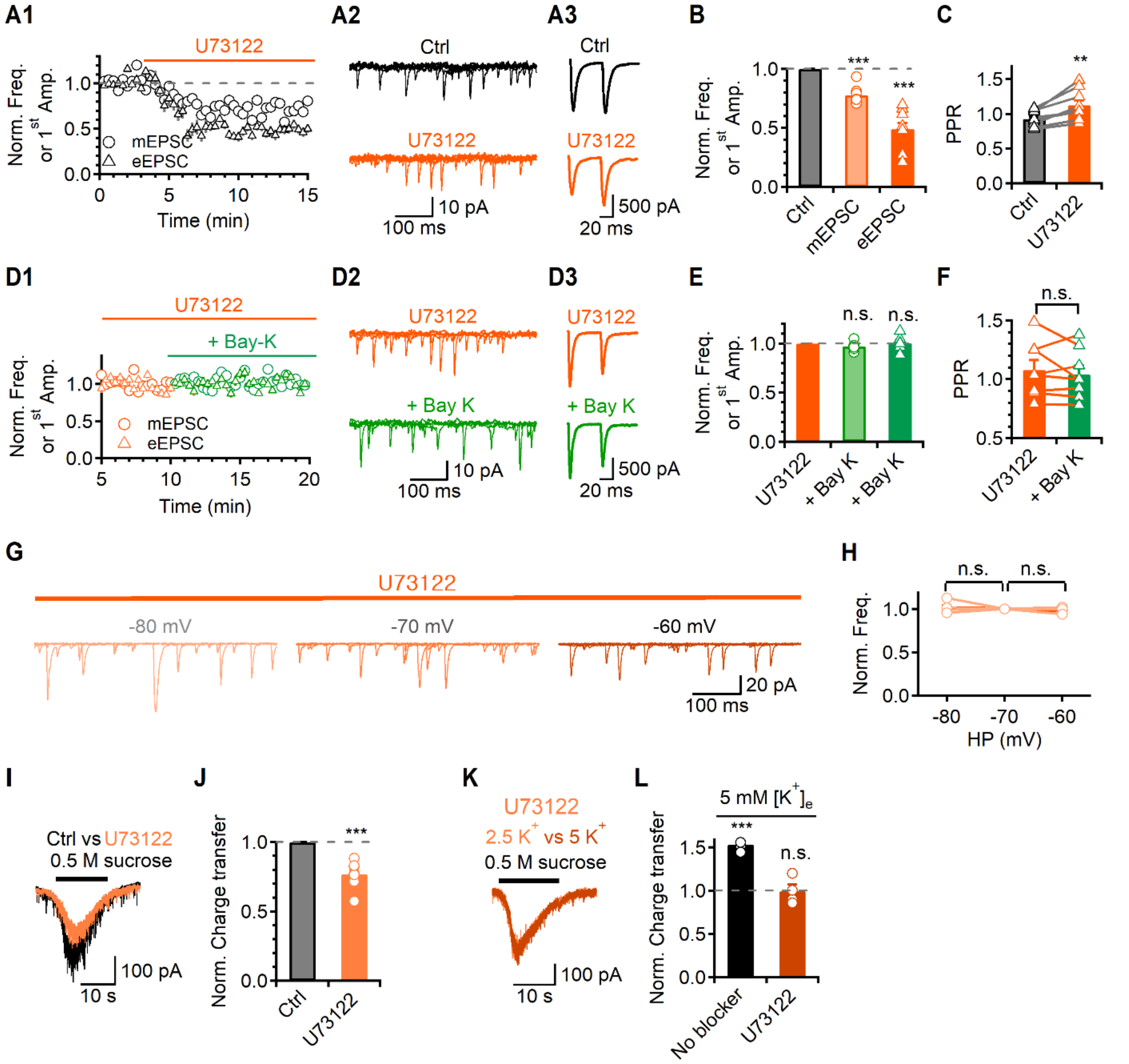


**Supplementary Fig. 9.** $V_m$- and LTCC-dependent regulation of release is mediated by PLC signaling.

(A) A1. An average time course of the normalized mEPSC frequency (circle) or the first eEPSC amplitude (triangle). The solid lines indicate the presence of U73122 (orange). A dashed grey line indicates the control level. A2, 3. Representative traces of mEPSCs (A2) or eEPSCs (A3) in control and U73122. (B) A bar graph of average values of the normalized mEPSC frequency (pale) and the first eEPSC amplitude (solid) in the presence of U73122, compared to control. (C) A bar graph of average values of the PPR in the presence of U73122. (D) D1. An average time course of the normalized mEPSC frequency (circle) and the normalized first eEPSC amplitude (triangle) in the presence of U73122 followed by Bay K. D2, 3. Representative traces of mEPSC (D2) or eEPSC (D3) in the presence of U73122 followed by Bay K, respectively. (E) A bar graph of average value of the normalized mEPSC frequency (pale green) and the first eEPSC amplitude (solid green) in the presence of U73122 followed by Bay K. (F) A bar graph of average values of the PPR. (G) Representative traces of mEPSC in each HP in the presence of U73122. (H) A graph showing the average value of the normalized mEPSC frequency in various HP in the presence of U73122, compared to -70 mV. (I) Representative traces of the hypertonic sucrose solution application in the presence of U73122. Top solid line in each trace indicates sucrose application periods. (J) A bar graph of the average values of the normalized charge transfer. (K) Representative traces of the hypertonic sucrose solution application for the comparison of 2.5 mM and 5 mM $[K^+]_e$ in the pretreatment U73122. (L) A bar graph of the average values of the normalized charge transfer in pretreatment of each blocker in 5 mM $[K^+]_e$, compared to 2.5 mM $[K^+]_e$. The individual raw values are described in table S1.

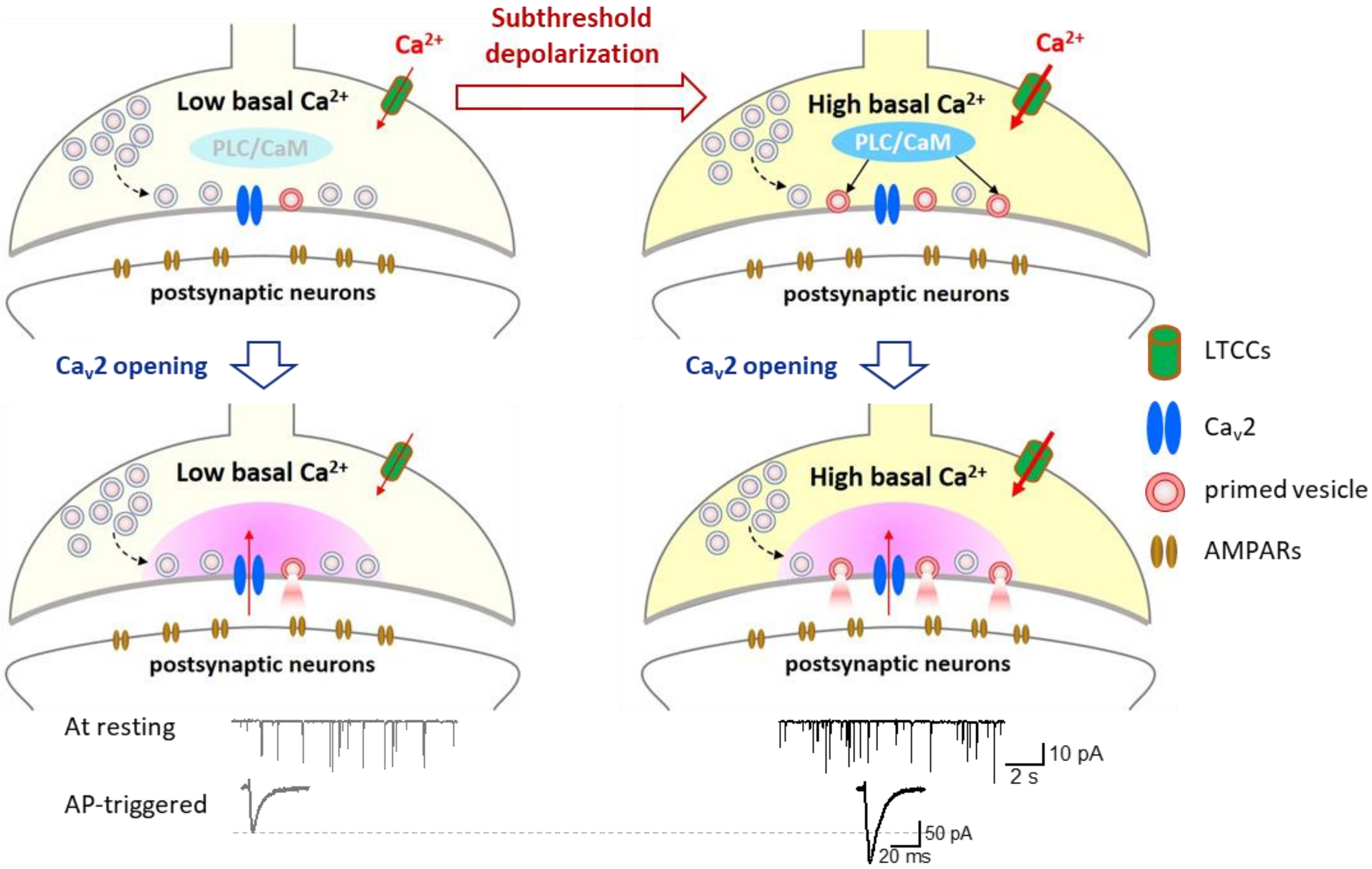


**Supplementary Fig. 10.** A summary cartoon to show the process involved in increased transmitter release by subthreshold depolarization.

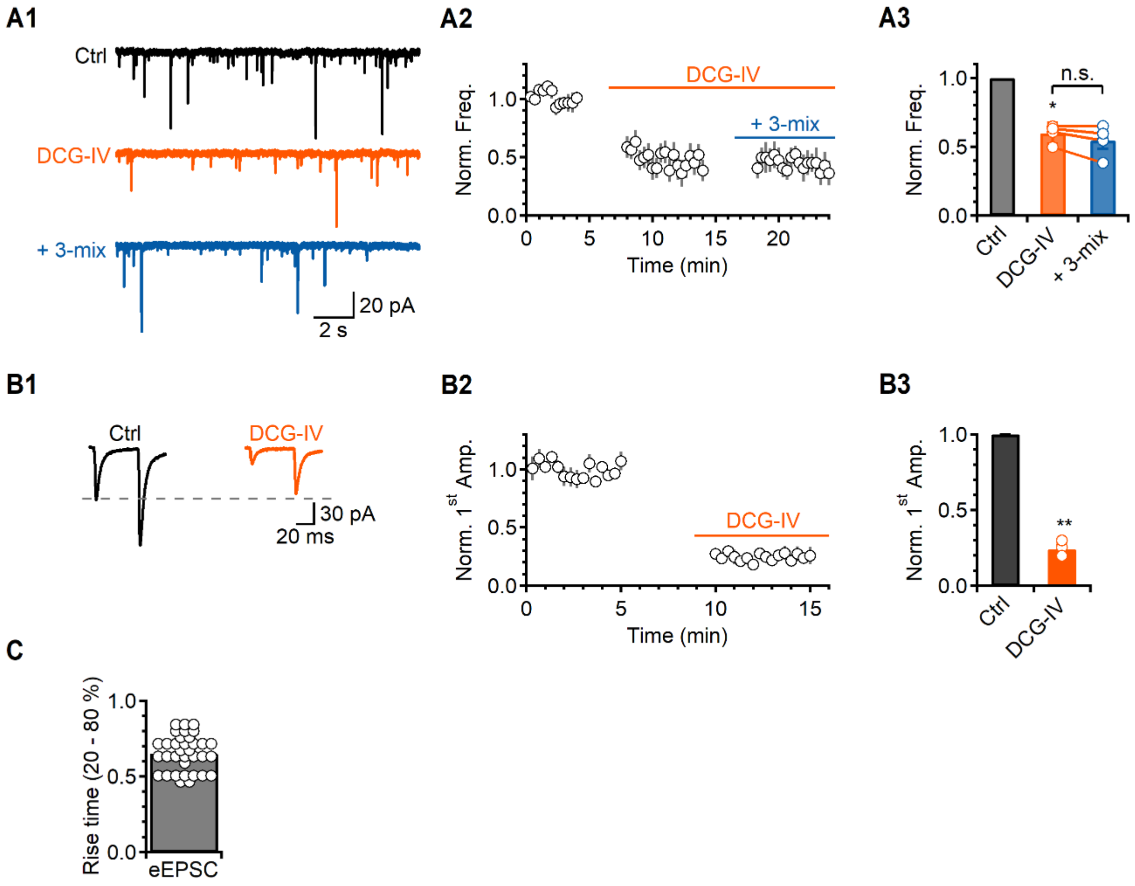


**Supplementary Fig. 11.** The effect of DCG-IV on mEPSC and eEPSC at hippocampal MF-CA3 synapse.

(A1) Representative traces of mEPSCs in control and the presence of DCG-IV (orange) followed by 3-mix (blue). (A2) Average time courses of the normalized mEPSC frequency. The upper solid line indicates the presence of DCG-IV and the lower solid line indicates the additional application of 3-mix. (A3) A bar graph of average values of the mEPSC frequency in presence of DCG-IV and additional treatment of 3-mix. (B1) Representative traces of eEPSCs in control and the presence of DCG-IV (orange). The grey dashed line indicates the control first eEPSC peak amplitude. (B2) Average time courses of the normalized first eEPSC amplitude. The upper solid line indicates the presence of DCG-IV. (B3) A bar graph of average values of the first eEPSC amplitude in presence of DCG-IV. (C) A bar graph of eEPSC rise time (20 - 80 %). The individual raw values are described in table S1.

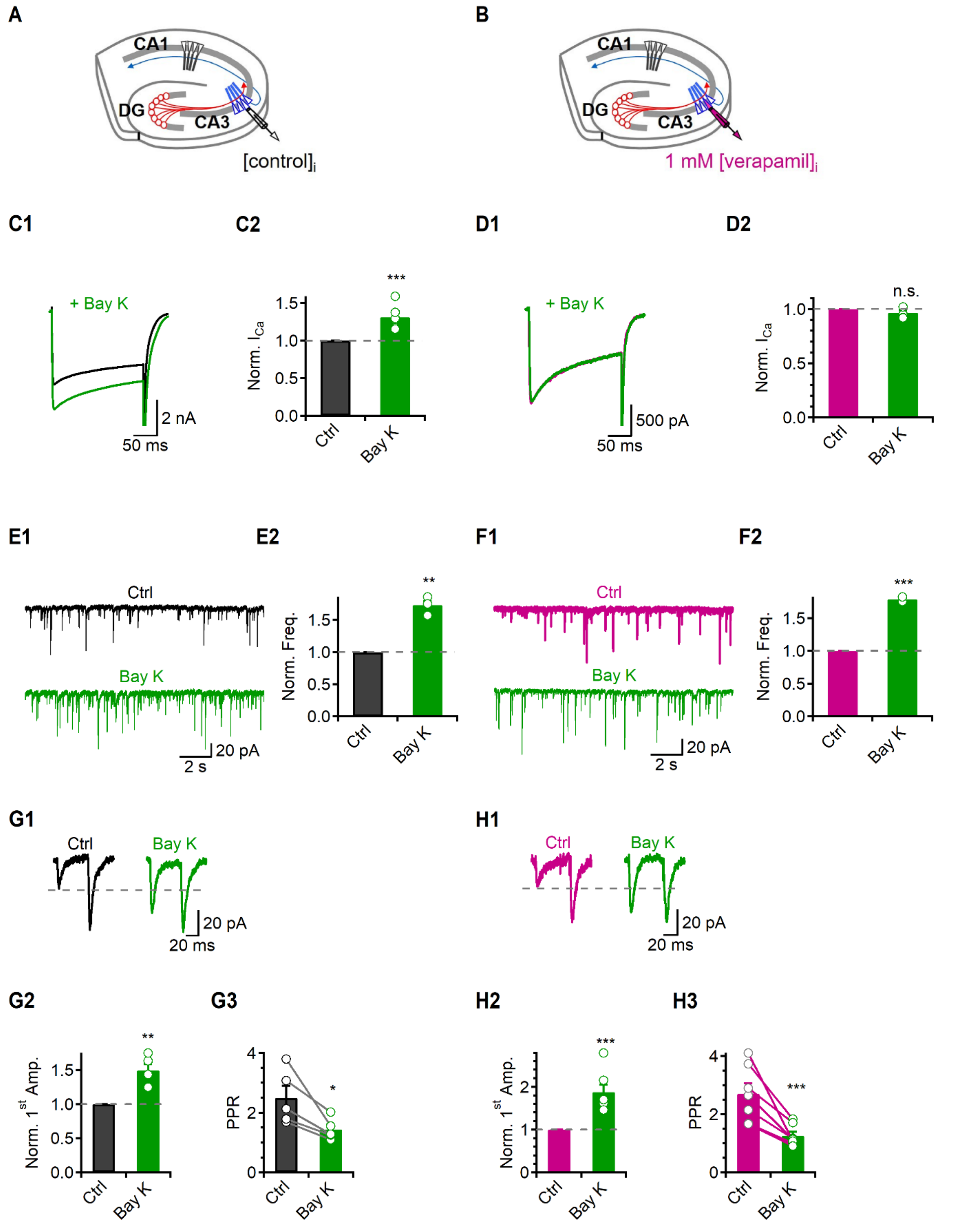


**Supplementary Fig. 12.** The effect of Bay K on $I_{Ca}$, mEPSC, and eEPSC at hippocampal MF-CA3 synapse using verapamil containing internal patch pipette.

(A, B) A schematic image of hippocampal slice in normal (A) and verapamil containing internal patch pipette solution (B) condition. Verapamil is an LTCCs blocker which binds to a pore-forming $Ca^{2+}$-conducting α1 subunit of LTCCs at high concentrations when intracellularly applied. (C1, D1) Representative trances of $I_{Ca}$ in normal (C1) and verapamil containing internal patch pipette solution (D1). (C2, D2) A bar graph of the average values of the normalized $I_{Ca}$ in presence of Bay K in normal

(C2) and verapamil condition (D2). In control condition, Bay K increased the somatic $I_{Ca}$ by 1.31 ± 0.07 fold (C2, *N* = 7), but $I_{Ca}$ was not changed by Bay K (D2, 0.97 ± 0.02, *N* = 5) in the verapamil containing patch pipette solution, confirming that verapamil could completely block the postsynaptic LTCCs. A dashed grey line indicates the control level. (E1, F1) Representative traces of mEPSCs in control and the presence of Bay K in normal (E1) and verapamil condition (F1). (E2, F2) A bar graph of the average values of the normalized mEPSC frequency in presence of Bay K in normal (E2) and verapamil condition (F2). (G1, H1) Representative traces of eEPSCs in control and the presence of Bay K in normal (G1) and verapamil condition (H1). The grey dashed line indicates the control first eEPSC peak amplitude. (G2, H2) A bar graph of the average values of the normalized first eEPSC amplitude in presence of Bay K in normal (G2) and verapamil condition (H2). (G3, H3) A bar graph of the average values of the PPR in presence of Bay K in normal (G3) and verapamil condition (H3). The individual raw values are described in table S1.

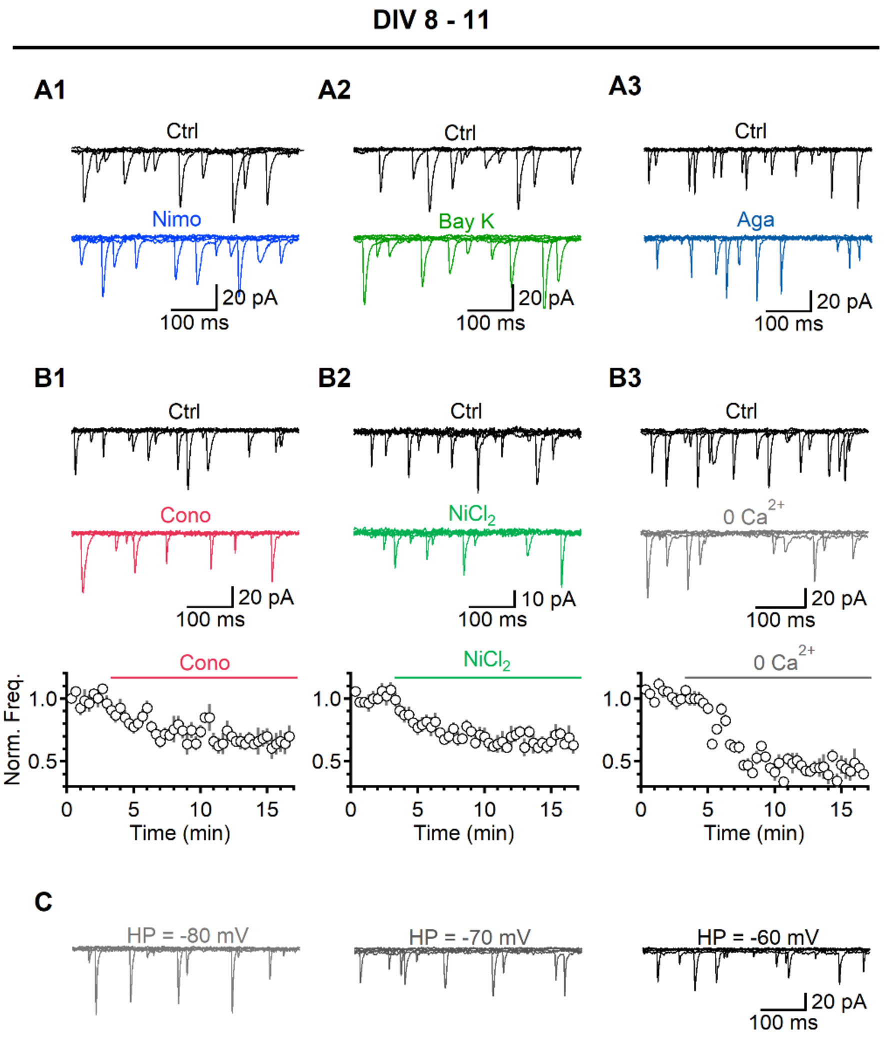


**Supplementary Fig. 13.** The effects of VGCCs or $Ca^{2+}$ removal on the spontaneous release in immature autaptic neurons.

(A) Representative traces of mEPSCs in different conditions (A1, Nimo; A2, Bay K; A3, Aga) for figure 6A. (B) Top. Representative traces of mEPSCs in control, Cono (B1, red), $NiCl_2$ (B2, green) and removal of $[Ca^{2+}]_e$ (B3). Five 500 ms-long mEPSC traces were overlaid. Bottom. Average time courses of the normalized mEPSC frequency. In each time course plot, the solid lines indicate the presence of each drug or removal of $[Ca^{2+}]_e$. (C) Representative traces of mEPSC frequency in each HP at 2.5 mM $[K^+]_e$. The data were normalized by the mean mEPSC frequency of control.

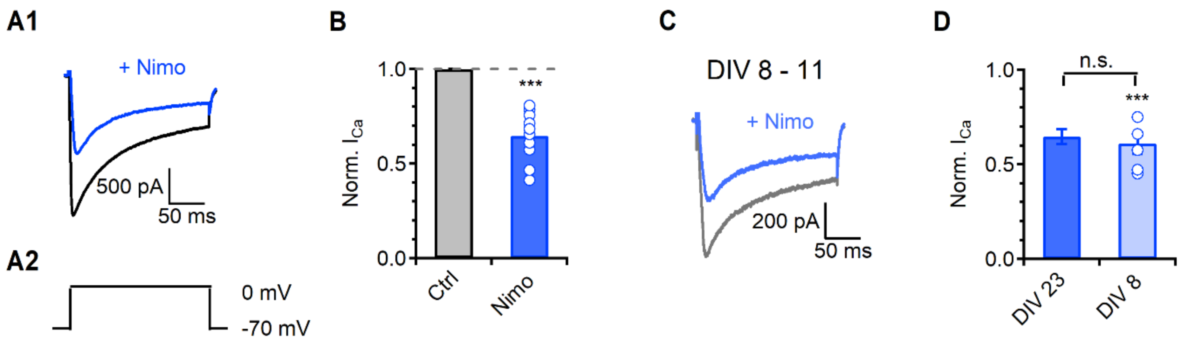


**Supplementary Fig. 14.** The effect of LTCCs on $I_{Ca}$.

(A1, C) Representative traces of $I_{Ca}$ applying Nimo (A) for comparison between DIV 23 ~ and DIV 8-11 (C). (A2) A pulse protocol for recording VGCC currents. The VGCC currents were activated by steps from -70 to 0 mV in bath solution containing 25 mM tetraethylammonium (TEA), 5 mM 4-AP, 1 μM TTX, 10 μM CNQX and 100 μM PTX. (B) A bar graph of the average values of the normalized $I_{Ca}$ in presence of Nimo. A dashed grey line indicates the control level. (D) A bar graph of the average values of the normalized $I_{Ca}$ in the presence of Nimo for comparison between DIV 23 ~ and DIV 8-11. The individual raw values are described in table S1.

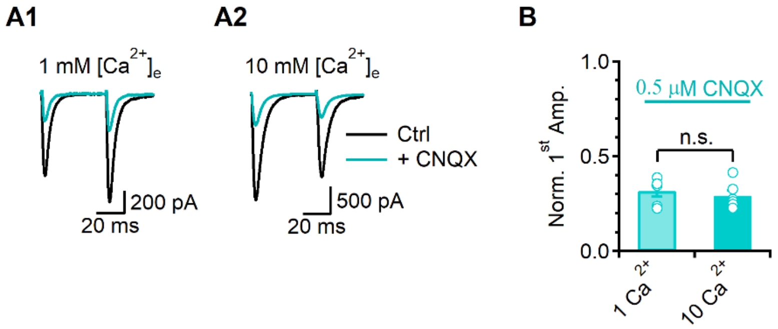


**Supplementary Fig. 15.** The effect of low dose of CNQX on eEPSC in 1 mM and 10 mM $[Ca^{2+}]_e$.

(A) Representative traces of eEPSCs at 1 mM (A1) and 10 mM $[Ca^{2+}]_e$ (A2) in the presence of 0.5 μM CNQX (cyan). (B) A bar graph of average values of the normalized first eEPSC amplitude in presence of CNQX. The individual raw values are described in table S1.

**Table S1.** Raw data from graphs.

| Figure | Graph | Contidion | Data | *N* | *p*-value | Analysis |
|---|---|---|---|---|---|---|
| **1B** | Norm. mEPSC Freq. | Nimo | 0.81 ± 0.01 | 17 | 8.38E-11 | Student's *t*-test |
| | | Bay K | 1.79 ± 0.11 | 7 | 0.001 | Student's *t*-test |
| **1C** | Norm. mEPSC Amp. | Nimo | 1.0 ± 0.02 | 17 | 0.822 | Student's *t*-test |
| | | Bay K | 0.99 ± 0.04 | 7 | 0.867 | Student's *t*-test |
| **1E** | Norm. 1st Amp. | Nimo | 0.7 ± 0.05 | 14 | 4.05E-05 | Student's *t*-test |
| | | Bay K | 1.82 ± 0.18 | 5 | 0.004 | Student's *t*-test |
| **1F** | PPR | Ctrl *vs* Nimo | 0.81 ± 0.07 *vs* 0.95 ± 0.08 | 7 | 0.014 | Student's paired *t*-test |
| | | Ctrl *vs* Bay K | 0.91 ± 0.09 *vs* 0.75 ± 0.06 | 5 | 0.046 | Student's paired *t*-test |
| **1H** | Norm. mEPSC Freq. | Nimo | 0.97 ± 0.03 | 5 | 0.277 | Student's *t*-test |
| | | Bay K | 1.02 ± 0.02 | 4 | 0.468 | Student's *t*-test |
| **1I** | Norm. eEPSC Amp. | Nimo | 0.96 ± 0.02 | 5 | 0.061 | Student's *t*-test |
| | | Bay K | 0.99 ± 0.05 | 4 | 0.904 | Student's *t*-test |
| **1J2** | Norm. mEPSC Freq. | Nimo | 1.01 ± 0.01 | 5 | 0.689 | Student's *t*-test |
| | | Bay K | 1.02 ± 0.02 | 5 | 0.254 | Student's *t*-test |
| **1K2** | Norm. 1st Amp. | Nimo | 0.94 ± 0.02 | 7 | 0.422 | Student's *t*-test |
| | | Bay K | 1.01 ± 0.03 | 7 | 0.802 | Student's *t*-test |

| | | | | | | |
|---|---|---|---|---|---|---|
| **2B** | Norm. mEPSC Freq. | -80 mV | 0.82 ± 0.02 | 12 | 2.86E-07 | Student's *t*-test |
| | | -60 mV | 1.49 ± 0.06 | 11 | 3.89E-05 | Student's *t*-test |
| | RMP vs Norm. Freq. | 1 $[K^+]_e$ | -85.33 ± 1.9 mV *vs* 0.72 ± 0.03 | 6 | | |
| | | 2.5 $[K^+]_e$ | -73.4 ± 1.37 mV *vs* 0.93 ± 0.04 | 7 | | |
| | | 5 $[K^+]_e$ | -54.67 ± 2.72 mV *vs* 1.43 ± 0.16 | 7 | | |
| **2D** | Norm. mEPSC Freq. | -80 mV | 0.99 ± 0.02 | 7 | 0.519 | Student's *t*-test |
| | | -60 mV | 0.99 ± 0.03 | 7 | 0.871 | Student's *t*-test |
| **2F** | Norm. mEPSC Freq. | -80 mV, Aga | 0.79 ± 0.04 | 4 | 0.010 | Student's paired *t*-test |
| | | -60 mV, Aga | 1.61 ± 0.05 | 7 | 2.2766.E-05 | Student's paired *t*-test |
| | | -80 mV, Nimo | 0.95 ± 0.03 | 4 | 0.179 | Student's paired *t*-test |
| | | -60 mV, Nimo | 1.02 ± 0.03 | 4 | 0.726 | Student's paired *t*-test |
| | | -80 mV, shCaV | 1.01 ± 0.02 | 4 | 0.708 | Student's paired *t*-test |
| | | -60 mV, $shCa_V$ | 1.01 ± 0.04 | 4 | 0.825 | Student's paired *t*-test |
| **2H** | Basal $Ca^{2+}$ level | 1 $[K^+]_e$, Ctrl | 0.961 ± 0.012 | 13 | 0.44 | Bonferroni test |
| | | 5 $[K^+]_e$, Ctrl | 1.025 ± 0.027 | 13 | | |
| | | 1 $[K^+]_e$, Nimo | 0.908 ± 0.015 | 15 | 0.36 | Bonferroni test (1 $K^+$ to 5 $K^+$) |
| | | 2.5 $[K^+]_e$, Nimo | 0.909 ± 0.027 | 11 | 0.012 | Student's *t*-test |
| | | 5 $[K^+]_e$, Nimo | 0.907 ± 0.025 | 15 | | |
| | | 1 $[K^+]_e$, Aga | 0.953 ± 0.011 | 12 | 0.006 | Bonferroni test (1 $K^+$ to 5 $K^+$) |

| | | | | | | |
|---|---|---|---|---|---|---|
| | | 2.5 $[K^+]_e$, Aga | 0.972 ± 0.013 | 12 | 0.057 | Student's *t*-test |
| | | 5 $[K^+]_e$, Aga | 0.991 ± 0.013 | 12 | | |
| **3C** | Peak release | 1 $[K^+]_e$, Ctrl | 0.73 ± 0.06 | 13 | 3.00.E-04 | Student's *t*-test |
| | | 5 $[K^+]_e$, Ctrl | 1.33 ± 0.07 | 13 | 4.00.E-06 | Student's *t*-test |
| | | 1 $K^+$ to 5 $K^+$, Ctrl | 1.82 ± 0.02 | | 1.12.E-07 | Bonferroni test (1 $K^+$ to 5 $K^+$) |
| | | 1 $[K^+]_e$, Nimo | 0.66 ± 0.05 | 15 | 0.079 | Student's *t*-test |
| | | 2.5 $[K^+]_e$, Nimo | 0.79 ± 0.02 | 11 | 3.98E-04 | Student's *t*-test |
| | | 5 $[K^+]_e$, Nimo | 0.88 ± 0.03 | 15 | 0.173 | Student's *t*-test |
| | | 1 $K^+$ to 5 $K^+$, Nimo | 1.33 ± 0.01 | | 5.75.E-05 | Bonferroni test (1 $K^+$ to 5 $K^+$) |
| | | 1 $[K^+]_e$, Aga | 0.56 ± 0.02 | 9 | 0.008 | Student's *t*-test |
| | | 2.5 $[K^+]_e$, Aga | 0.64 ± 0.02 | 9 | 4.00E-07 | Student's *t*-test |
| | | 5 $[K^+]_e$, Aga | 0.77 ± 0.03 | 9 | 1.00E-05 | Student's *t*-test |
| | | 1 $K^+$ to 5 $K^+$, Aga | 1.36 ± 0.02 | | 7.95E-09 | Bonferroni test (1 $K^+$ to 5 $K^+$) |
| **3F** | Peak $Ca^{2+}$ | 1 $[K^+]_e$, Ctrl | 0.85 ± 0.02 | 13 | 3.00.E-04 | Student's *t*-test |
| | | 5 $[K^+]_e$, Ctrl | 1.11 ± 0.02 | 13 | 4.00.E-06 | Student's *t*-test |
| | | 1 $[K^+]_e$, Nimo | 0.76 ± 0.02 | 15 | 8.00.E-07 | Student's *t*-test |
| | | 2.5 $[K^+]_e$, Nimo | 0.88 ± 0.02 | 11 | 3.98E-04 | Student's *t*-test |
| | | 5 $[K^+]_e$, Nimo | 0.97 ± 0.02 | 15 | 9.00.E-05 | Student's *t*-test |
| | | 1 $[K^+]_e$, Aga | 0.7192 ± 0.016 | 12 | 4.00.E-05 | Student's *t*-test |

| | | | | | | |
|---|---|---|---|---|---|---|
| | | 2.5 $[K^+]_e$, Aga | 0.801 ± 0.02 | 12 | 9.00E-07 | Student's *t*-test |
| | | 5 $[K^+]_e$, Aga | 0.848 ± 0.019 | 12 | 0.01 | Student's *t*-test |
| **4B** | Norm. mEPSC Freq. | CaM-ip | 0.77 ± 0.04 | 11 | 7.00.E-03 | Student's *t*-test |
| | Norm. 1$^{st}$ Amp. | | 0.77 ± 0.06 | 13 | 1.70.E-04 | Student's *t*-test |
| **4C** | PPR | Ctrl *vs* CaM-ip | 1.02 ± 0.07 *vs* 1.15 ± 0.07 | 13 | 5.33.E-04 | Student's paired *t*-test |
| **4E** | Norm. mEPSC Freq. | Nimo | 0.96 ± 0.03 | 4 | 0.283 | Student's *t*-test |
| | | Bay K | 0.98 ± 0.02 | 4 | 0.390 | Student's *t*-test |
| **4G** | Norm. 1$^{st}$ Amp. | Nimo | 0.95 ± 0.05 | 4 | 0.362 | Student's *t*-test |
| | | Bay K | 0.99 ± 0.01 | 4 | 0.395 | Student's *t*-test |
| **4H** | PPR | Ctrl *vs* Nimo | 1.16 ± 0.12 *vs* 1.19 ± 0.11 | 4 | 0.703 | Student's paired *t*-test |
| | | Ctrl *vs* Bay K | 0.98 ± 0.09 *vs* 0.95 ± 0.09 | 4 | 0.282 | Student's paired *t*-test |
| **4J** | Norm. mEPSC Freq. | -80 mV | 1.01 ± 0.03 | 4 | 0.755 | Student's *t*-test |
| | | -60 mV | 1.01 ± 0.03 | 4 | 0.846 | Student's *t*-test |
| **4L** | Norm. charge transfer | Nimo | 0.73 ± 0.03 | 8 | 3.17E-06 | Student's *t*-test |
| | | CaM-ip | 0.61 ± 0.05 | 6 | 6.23E-04 | Student's *t*-test |
| **4N** | Norm. charge transfer | 5 $[K^+]_e$, No blocker | 1.53 ± 0.19 | 4 | 6.34E-04 | Student's *t*-test |
| | | 5 $[K^+]_e$, Nimo | 1.0 ± 0.05 | 6 | 0.955 | Student's *t*-test |
| | | 5 $[K^+]_e$, CaM-ip | 1.09 ± 0.05 | 4 | 0.171 | Student's *t*-test |
| **5B** | Norm. mEPSC Freq. | Nimo | 0.72 ± 0.04 | 15 | 5.00E-07 | Student's *t*-test |

| | | | | | | |
|---|---|---|---|---|---|---|
| | | CMZ | 0.55 ± 0.05 | 12 | 2.36E-06 | Student's *t*-test |
| | | U73122 | 0.78 ± 0.02 | 4 | 0.007 | Student's *t*-test |
| **5D2** | Norm. mEPSC Freq. | 5 $[K^+]_e$, No blocker | 1.76 ± 0.19 | 7 | 0.008 | Student's *t*-test |
| | | 5 $[K^+]_e$, Nimo | 1.06 ± 0.07 | 10 | 0.397 | Student's *t*-test |
| | | 5 $[K^+]_e$, CMZ | 1.04 ± 0.05 | 7 | 0.512 | Student's *t*-test |
| | | 5 $[K^+]_e$, U73122 | 1.01 ± 0.06 | 5 | 0.888 | Student's *t*-test |
| **5I1** | Norm. 1$^{st}$ Amp. | Nimo | 0.6 ± 0.04 | 4 | 0.002 | Student's *t*-test |
| | | CMZ | 0.62 ± 0.07 | 8 | 0.003 | Student's *t*-test |
| | | U73122 | 0.61 ± 0.06 | 9 | 0.016 | Student's *t*-test |
| **5I2** | Norm. 1$^{st}$ Amp. | 5 $[K^+]_e$, No blocker | 1.86 ± 0.12 | 7 | 2.20E-06 | Student's *t*-test |
| | | 5 $[K^+]_e$, Nimo | 0.99 ± 0.02 | 4 | 0.649 | Student's *t*-test |
| | | 5 $[K^+]_e$, CMZ | 0.96 ± 0.1 | 4 | 0.483 | Student's *t*-test |
| | | 5 $[K^+]_e$, U73122 | 0.99 ± 0.1 | 4 | 0.941 | Student's *t*-test |
| **5J** | PPR | Ctrl *vs* Nimo | 1.5 ± 0.07 *vs* 2.56 ± 0.12 | 4 | 0.024 | Student's paired *t*-test |
| | | Ctrl *vs* CMZ | 1.28 ± 0.07 *vs* 2.01 ± 0.25 | 4 | 0.050 | Student's paired *t*-test |
| | | Ctrl *vs* U73122 | 1.6 ± 0.24 *vs* 2.11 ± 0.14 | 4 | 0.006 | Student's paired *t*-test |
| | | Ctrl *vs* 5 $K^+$ | 2.0 ± 0.09 *vs* 1.32 ± 0.11 | 7 | 4.52E-04 | Student's paired *t*-test |
| **6B** | Norm. mEPSC Freq. | Nimo | 0.99 ± 0.02 | 6 | 0.669 | Student's *t*-test |
| | | Bay K | 1.03 ± 0.04 | 4 | 0.326 | Student's *t*-test |

| | | | | | | |
|---|---|---|---|---|---|---|
| | | Aga | 0.63 ± 0.04 | 5 | 2.29.E-04 | Student's *t*-test |
| | | Cono | 0.63 ± 0.04 | 7 | 3.21E-06 | Student's *t*-test |
| | | $NiCl_2$ | 0.7 ± 0.03 | 5 | 1.86E-06 | Student's *t*-test |
| **6C** | Norm. mEPSC Freq. | -80 mV | 0.99 ± 0.05 | 5 | 0.393 | Student's *t*-test |
| | | -60 mV | 1.02 ± 0.01 | 5 | 0.191 | Student's *t*-test |
| **6E** | Mander's colocalization coefficients | P/Q | 0.46 ± 0.04 | 5 | 4.81E-05 | Student's *t*-test |
| | | L | 0.23 ± 0.02 | 5 | | |
| **6G** | Mander's colocalization coefficients | DIV 8-11 | 0.08 ± 0.02 | 5 | 0.004 | Student's *t*-test |
| | | DIV 23 ~ | 0.22 ± 0.02 | 5 | | |
| **S1** | Decay time constant | EPSC | 6.9 ± 0.2 | 94 | | |
| | | IPSC | 47.4 ± 4.6 | 21 | | |
| **S4C** | Norm. mEPSC Freq. | Aga | 0.78 ± 0.03 | 5 | 0.003 | Student's *t*-test |
| | | Cono | 0.79 ± 0.02 | 4 | 5.29E-06 | Student's *t*-test |
| | | $NiCl_2$ | 0.76 ± 0.03 | 6 | 0.006 | Student's *t*-test |
| | | 0 $Ca^{2+}$ | 0.42 ± 0.02 | 4 | 3.17E-06 | Student's *t*-test |
| **S4D** | Relative contribution | P/Q | 0.53 ± 0.03 *vs* 0.42 ± 0.06 | 24 (0.1 EGTA) | 0.714 | |
| | | N | 0.49 ± 0.02 *vs* 0.39 ± 0.03 | 18 (0.1 EGTA) | 0.138 | |
| | | R | 0.42 ± 0.03 *vs* 0.45 ± 0.05 | 15 (0.1 EGTA) | 0.571 | |
| **S5A3** | Norm. mEPSC Freq. | 3-mix | 0.47 ± 0.01 | 6 | 5.74E-07 | Student's *t*-test |

| | | | | | | |
|---|---|---|---|---|---|---|
| | | + Bay K | 0.47 ± 0.01 | 6 | 0.438 | Student's paired *t*-test |
| **S5B3** | Norm. 1st Amp. | 3-mix | 0.039 ± 0.006 | 7 | 5.08.E-07 | Student's *t*-test |
| | | + Bay K | 0.037 ± 0.005 | 7 | 0.149 | Student's paired *t*-test |
| **S6** | mEPSC Amp. | -90 mV | 18.06 ± 0.88 pA | 11 | | |
| | | -80 mV | 17.66 ± 0.68 pA | 12 | | |
| | | -70 mV | 15.98 ± 0.54 pA | 12 | | |
| | | -60 mV | 14.2 ± 0.56 pA | 11 | | |
| **S7B** | Norm. mEPSC Freq. | -80 mV, 40 μM $NiCl_2$ | 0.79 ± 0.09 | 5 | 0.037 | Student's *t*-test |
| | | -70 mV, 40 μM $NiCl_2$ | 1.0 ± 0.01 | 5 | 0.731 | Student's *t*-test |
| | | -60 mV, 40 μM $NiCl_2$ | 1.37 ± 0.03 | 5 | 2.82E-04 | Student's *t*-test |
| | | | | | 0.528 | Student's paired *t*-test (*vs* -60 mV, Ctrl) |
| **S8C** | Norm. mEPSC Freq. | CaM-ip Scramble | 0.98 ± 0.02 | 5 | 0.542 | Student's *t*-test |
| | Norm. 1st Amp. | CaM-ip Scramble | 0.99 ± 0.03 | 5 | 0.792 | Student's *t*-test |
| **S9B** | Norm. mEPSC Freq. | U73122 | 0.77 ± 0.03 | 9 | 5.48E-05 | Student's *t*-test |
| | Norm. 1st Amp. | | 0.49 ± 0.05 | 10 | 1.39E-05 | Student's *t*-test |
| **S9C** | PPR | Ctrl *vs* U73122 | 0.93 ± 0.04 *vs* 1.13 ± 0.07 | 10 | 0.001 | Student's paired *t*-test |
| **S9E** | Norm. mEPSC Freq. | Bay K | 0.96 ± 0.02 | 6 | 0.269 | Student's *t*-test |
| | Norm. 1st Amp. | | 1.0 ± 0.02 | 9 | 0.916 | Student's *t*-test |
| **S9F** | PPR | Ctrl *vs* Bay K | 1.08 ± 0.08 *vs* 1.04 ± 0.07 | 9 | 0.436 | Student's paired *t*-test |

| | | | | | | |
|---|---|---|---|---|---|---|
| **S9H** | Norm. mEPSC Freq. | -80 mV | 1.01 ± 0.03 | 6 | 0.644 | Student's *t*-test |
| | | -60 mV | 0.98 ± 0.02 | 6 | 0.341 | Student's *t*-test |
| **S9J** | Norm. charge transfer | U73122 | 0.77 ± 0.04 | 7 | 8.28E-04 | Student's *t*-test |
| **S9L** | | 5 $[K^+]_e$, U73122 | 0.99 ± 0.08 | 4 | 0.895 | Student's *t*-test |
| **S11A3** | Norm. mEPSC Freq. | DCG-IV | 0.59 ± 0.08 | 4 | 0.013 | Student's *t*-test |
| | | + 3-mix | 0.54 ± 0.06 | 4 | 0.458 | Student's paired *t*-test |
| **S11B3** | Norm. 1st Amp. | DCG-IV | 0.24 ± 0.03 | 3 | 0.0015 | Student's *t*-test |
| **S11C** | Rise time | eEPSC | 0.65 ± 0.018 | 34 | | |
| **S12C2** | Norm. $I_{ca}$ | Bay K | 1.31 ± 0.065 | 6 | 0.005 | Student's *t*-test |
| **S12D2** | Norm. $I_{ca}$ | Bay K | 0.97 ± 0.022 | 5 | 0.202 | Student's *t*-test |
| **S12E2** | Norm. mEPSC Freq. | Bay K | 1.73 ± 0.014 | 4 | 0.001 | Student's *t*-test |
| **S12F2** | Norm. mEPSC Freq. | Bay K | 1.79 ± 0.0012 | 3 | 6.489.E-04 | Student's *t*-test |
| **S12G2** | Norm. 1st Amp. | Bay K | 1.5 ± 0.04 | 5 | 0.005 | Student's *t*-test |
| **S12G3** | PPR | Ctrl *vs* Bay K | 2.5 ± 0.41 *vs* 1.44 ± 0.16 | | 0.028 | Student's paired *t*-test |
| **S12H2** | Norm. 1st Amp. | Bay K | 1.87 ± 0.17 | 6 | 0.002 | Student's *t*-test |
| **S12H3** | PPR | Ctrl *vs* Bay K | 2.7 ± 0.36 *vs* 1.26 ± 0.14 | | 0.004 | Student's paired *t*-test |
| **S14B** | Norm. $I_{ca}$ | Nimo | 0.65 ± 0.04 | 11 | 2.95E-06 | Student's *t*-test |
| **S14D** | Norm. $I_{ca}$ | DIV 8-11 | 0.61 ± 0.06 | 6 | 7.77.E-04 | Student's *t*-test |
| | | DIV 23 ~ | 0.65 ± 0.04 | 11 | 0.577 | Student's paired *t*-test |

| | | | | | | |
|---|---|---|---|---|---|---|
| **S15B** | Norm. 1st Amp. | 1 mM $[Ca^{2+}]_e$ | 0.31 ± 0.03 | 6 | 2.043E-06 | Student's *t*-test |
| | | 10 mM $[Ca^{2+}]_e$ | 0.29 ± 0.03 | 6 | 0.558 | Student's paired *t*-test |
